# SPECIFIC ECTODERMAL ENHANCERS CONTROL THE EXPRESSION OF *Hoxc* GENES IN DEVELOPING MAMMALIAN INTEGUMENTS

**DOI:** 10.1101/2020.06.10.143677

**Authors:** Marc Fernandez-Guerrero, Nayuta Yakushiji-Kaminatsui, Lucille Lopez-Delisle, Sofía Zdral, Fabrice Darbellay, Rocío Perez-Gomez, Christopher Chase Bolt, Manuel A. Sanchez-Martin, Denis Duboule, Maria A. Ros

## Abstract

Vertebrate *Hox* genes are key players in the establishment of structures during the development of the main body axis. Subsequently, they play important roles either in organizing secondary axial structures such as the appendages, or during homeostasis in postnatal stages and adulthood. Here we set up to analyze their elusive function in the ectodermal compartment, using the mouse limb bud as a model. We report that the *HoxC* gene cluster was globally co-opted to be transcribed in the distal limb ectoderm, where it is activated following the rule of temporal colinearity. These ectodermal cells subsequently produce various keratinized organs such as nails or claws. Accordingly, deletion of the *HoxC* cluster led to mice lacking nails (anonychia) and also hairs (alopecia), a condition stronger than the previously reported loss of function of *Hoxc13*, which is the causative gene of the ectodermal dysplasia 9 (ECTD9) in human patients. We further identified two ectodermal, mammalian-specific enhancers located upstream of the *HoxC* gene cluster, which act synergistically to regulate *Hoxc* gene expression in the hair and nail ectodermal organs. Deletion of these regulatory elements alone or in combination revealed a strong quantitative component in the regulation of *Hoxc* genes in the ectoderm, suggesting that these two enhancers may have evolved along with mammals to provide the level of HOXC proteins necessary for the full development of hairs and nails.

**Significance Statement:** In this study, we report a unique and necessary function for the *HoxC* gene cluster in the development of some ectodermal organs, as illustrated both by the hair and nail phenotype displayed by mice lacking the *Hoxc13* function and by the congenital anonychia (absence of nails) in full *HoxC* cluster mutants. We show that *Hoxc* genes are activated in a colinear manner in the embryonic limb ectoderm and are subsequently transcribed in developing nails and hairs. We identify two mammalian-specific enhancers located upstream of the *HoxC* cluster with and exclusive ectodermal specificity. Individual or combined enhancer deletions suggest that they act in combination to raise the transcription level of several *Hoxc* genes during hairs and nails development.

## Introduction

In most bilateria, genes members of the *Hox* family of transcription factors are important for the proper organization of structures along the main body axis, during early development. On top of this major task, vertebrate *Hox* genes are also necessary for the proper morphogenesis of a variety of secondary structures such as the appendages or the external genitalia (1). In amniotes, *Hox* genes are organized in four clusters (*HoxA*, *B*, *C*, and *D*), all of them being involved during the development of the trunk axis. Subsequently however, sub-groups of gene clusters display some specificities regarding where and when to act in a global functional manner, usually with functional redundancy. For example, both the *HoxA* and *HoxD* clusters are essential for the organization of the tetrapod limb morphology (2–7), while the deletion of both *HoxA* and *HoxB* clusters leads to severe cardiac malformations, not detected in any of the single deletion mutants. Likewise, deletions of both the *HoxA* and *HoxC* clusters induced a complete renal aplasia, which was not detected with either deletion alone (8).

Such a functional redundancy between *Hox* clusters as a whole can be explained by their patterns of duplication during the phylogenetic history of vertebrates, with particular functional traits being conserved in all or some gene clusters after duplication (8). In this context, *Hox* gene function associated with recent vertebrate features, i.e. features that must have appeared late after the two rounds of duplication, can be expected to involve single gene clusters and thus to show less functional redundancy. Examples of this situation may be found either in the function of some *Hoxa* genes in uterine physiology (9–11) or in the apparent specialization of some *Hoxc* genes for epidermal derivatives (12, 13), while other clusters apparently do not play any detectable function there.

Indeed, while the single deletion of the *HoxC* gene cluster did not reveal any gross phenotype (14), several studies have suggested that *Hoxc* genes are involved in the development of ectodermal organs (12, 15). The fact that such organs usually become fully developed after birth explains why the full *HoxC* cluster deletion, which is lethal at birth because of respiratory problems (14), did not unmask these functional contributions. However, both expression and inactivation studies have revealed potential contributions, in particular at late fetal stages as well as during postnatal development and even adulthood (12, 16, 17). In such instances, some *Hoxc* genes display complex patterns of expression in relation to the differentiation of epidermal organs such as the hairs and nails (12, 15, 18, 19), two ectodermal derivatives which form through epithelial– mesenchymal interactions between the surface ectoderm and the underlying mesoderm, the latter being responsible for the development of site-specific skin derivatives (17, 20).

The morphogenesis of both hairs and nails begins with the induction of the corresponding placodes, which are ectodermal thickenings, by a subjacent dermal condensate (21). Once the placode forms, signaling events between the ectodermal and mesodermal components drive proliferation of the ectoderm to produce the hair peg or the nail matrix and eventually lead to a mature hair follicle or nail organ. Both the morphogenetic steps and the signaling molecules involved are remarkably similar during the development of these two integumentary structures (reviewed in 22, 23) and there is evidence suggesting that *Hoxc* genes may participate in these processes. This is mostly supported by the inactivation of *Hoxc13*, which induced specific hair and nail defects in adult homozygous mutant animals (12, 13). These mice showed long and fragile nails, they lacked vibrissae at birth and subsequently displayed a nude appearance with no external pelage. Although the morphogenesis of hair follicles appeared normal, hair differentiation was defective with very fragile shafts that broke as soon as they emerged through the skin surface, leading to alopecia. Both the hair and claw defects were associated with defective keratin gene expression, as *Hoxc13* was reported to be an important regulator of various keratin genes (15).

In order to evaluate the regulatory strategy at work at the *HoxC* locus, i.e. to assay first how many *Hoxc* genes are involved into the development of such epidermal organs and, secondly, whether a single global ‘epidermal regulatory modality’ may control their expression there in a coordinated manner, we selected the developing mouse limb bud as a privileged model system allowing to micro-dissect out the ectodermal component from where nails and hairs subsequently arise. We used RNA-seq on this isolated ectodermal jacket during both the early patterning phase and also the subsequent phase of tissue-specific differentiation. Here we show that *Hoxc* genes, unlike both *Hoxa* and *Hoxd* genes, are specifically expressed in a colinear manner in the limb bud ectoderm. At later fetal stages, expression becomes localized in the developing nail region and hair follicles (HF). Accordingly, we report that mice lacking the *HoxC* cluster suffer from anonychia (absence of nail or claw) at birth. We show that this global transcriptional control is in large part associated with two functionally redundant *cis*-regulatory regions, for only the deletion of both enhancers together led to a phenotype involving a disheveled fur and hypoplastic nails. Finally, the removal of both enhancers in a weakened *HoxC* genetic background caused a striking cyclic alopecia. We conclude that the *HoxC* cluster was evolutionary co-opted for exerting an essential function during the development of the integument. As reported for other *Hox* clusters in different contexts, this co-option was at least partly achieved through the emergence of remote enhancers with ectodermal specificities, taking control of series of *Hoxc* genes in a colinear manner.

## Results

### Colinear expression of *Hoxc* genes in the limb bud ectoderm

Initially, we set out to study in details how *Hoxc* genes are regulated in the developing limb by carefully isolating the ectodermal layer from the mesoderm component. To this aim, we generated RNA-seq datasets of both micro-dissected mouse distal forelimb mesenchymal progenitors and the overlying ectodermal cells at four consecutive developmental stages. We selected E9.5, E10.5, E11.5 and E12.5 limb buds because they illustrate a wide range of processes, from the early emergence of the limb bud to the specification of the digits. Due to the progressive proximo-distal development of the limb bud, our analysis was restricted to the distal 150 microns, which was considered a good approximation of the progress zone (24–26). As consequence, we micro-dissected a 150 microns thick band of the distal limb bud and the mesoderm and ectoderm components were separated by mild trypsin digestion and processed separately (Fig. S1A, see Materials & Methods). Two biological replicates were collected per stage and each pair of ectoderm and mesoderm replicates was obtained from the same batch of embryos.

Because a few mesoderm cells could occasionally remain attached to the ectoderm during dissection, potentially contaminating the ectodermal samples, we evaluated the purity of samples by analysing the expression of genes known to be expressed exclusively in the ectoderm or mesoderm components of limb buds. *Fgf8, Sp6, Trp63, Perp, Krt14, Sp8* and *Wnt7a* were selected as ectoderm specific genes, whereas *Prrx1, Tbx5, Hoxa9, Hoxd10, Fgf10, Pecam1* and *Cdh5* were selected as mesoderm specific genes (1, 27–31). A heatmap visualization of the transcription profiles of these genes across samples showed that the separation of the ectoderm and the mesoderm was clean, with a negligible level of contamination between both limb components (Fig. S1B), except in replicate 1 of the E11.5 ectoderm samples where a slight contamination with mesodermal cells was scored (see Fig. 1A).

**Figure 1.**
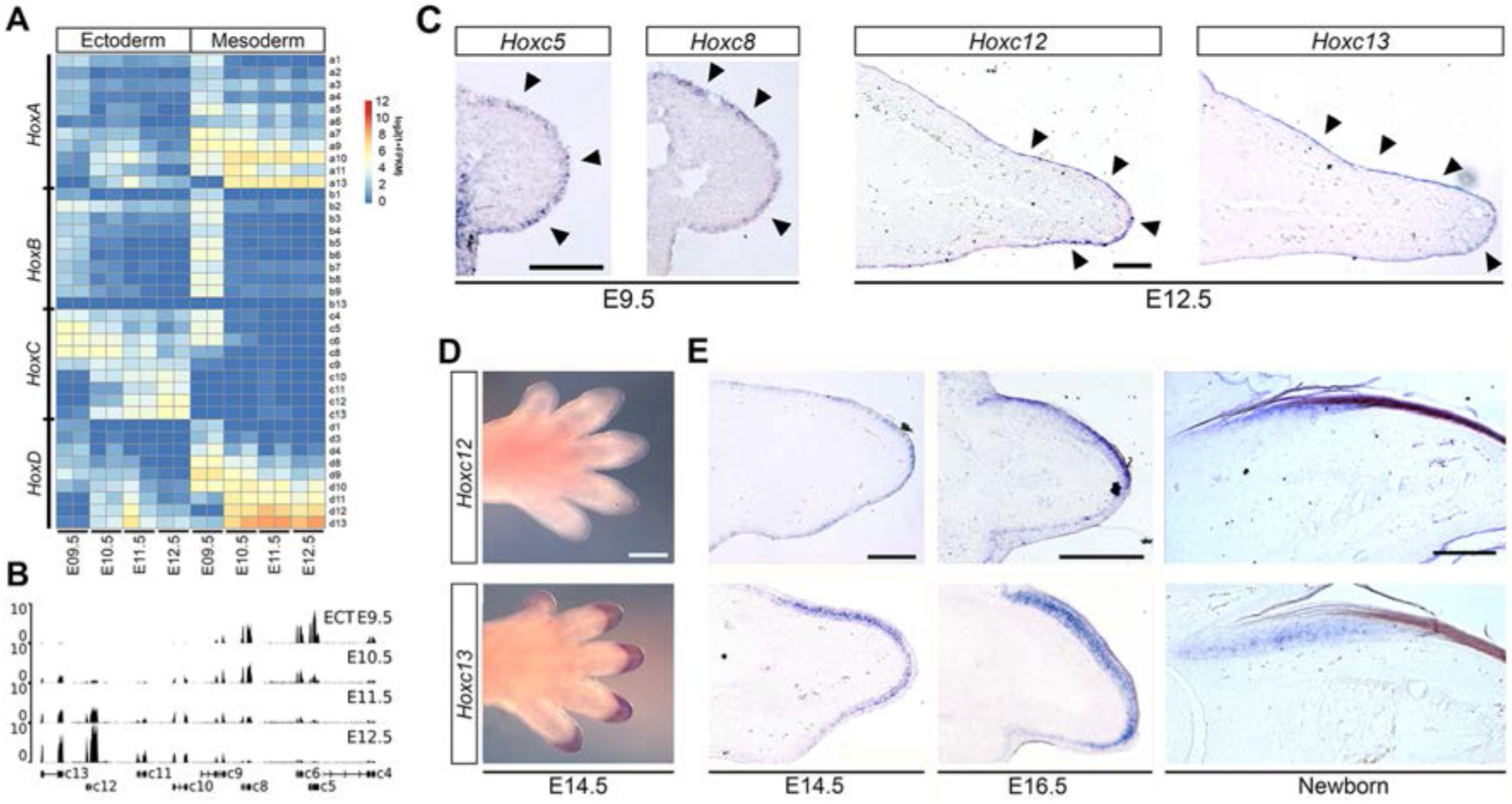
Colinear activation of *Hoxc* genes in the limb ectoderm. **A**) Heatmap showing the log2(1+FPKM) values of all *Hox* genes across the samples. *Hoxc* genes are activated in the ectoderm following a temporal sequence. **B**) Expression profiles of *HoxC* cluster genes in the four stages analyzed. Data were normalized to the million uniquely mapped reads and mean of duplicates are presented. **C**) mRNA *in situ* hybridization in tissue sections showing expression of *Hoxc5* and *Hoxc8* in the early E9.5 limb bud ectoderm and of *Hoxc12* and *Hoxc13* in the late E12.5 limb bud ectoderm. Arrowheads indicate the ectodermal restriction of expression. Scale bars 100μm. **D**) Whole mount *in situ* hybridization showing restriction of *Hoxc12* and *Hoxc13* expression to the forelimb digit tips at E14.5. Scale bar 500μm. **E)** *Hoxc12* and *Hoxc13* transcripts become progressive confined to the dorsal ectoderm over the last phalanx (E16.5) and finally to the nail matrix in newborn. For components of the nail organ please see Fig. 2C. Scale bars are 200μm. In all sections dorsal is up and distal to the right. For components of the nail organ please refer to Fig. 2C.

To evaluate the proximity between samples, we performed a principal component analysis (PCA). When samples were plotted along the first two components and in two dimensions, the first dimension (PC1), accounting for 75 percent of the variance, strongly separated the samples according to the ectodermal and mesodermal nature of the tissue. The second dimension (PC2), accounting for 12 percent of the variance, separated the samples according to the developmental stage (Fig. S1C). Hierarchical cluster analysis (HCA) of the 16 samples showed a strong clustering of samples by tissue and between consecutive stages (Fig. S1D). Both the PCA and the HCA showed a high consistency between replicates.

We then looked at the tissue specific gene expression dynamics of all *Hox* family members and generated a heatmap using their normalized expression values (log2(1+FPKM)). While this plot expectedly highlighted the known temporal colinear activation of *Hoxa* and *Hoxd* genes in the limb mesoderm (Fig. 1A), it revealed a previously overlooked distinctive expression of *Hoxc* genes in the limb ectoderm, with a comparable progressive temporal activation, starting with 3’-located genes and with 5’-located *Hoxc12* and *Hoxc13* gene expression being maximum at the last stage examined (E12.5) (Fig. 1A). The visual inspection of coverage from RNA-seq across the *HoxC* cluster confirmed a transition from an initial higher expression of *Hoxc5*, *Hoxc6* and *Hoxc8* (E9.5 and E10.5) to a higher expression of *Hoxc12* and *Hoxc13* at later stages (E11.5 and E12.5) (Fig. 1B). Genes with an intermediate position in the cluster such as *Hoxc9*, *Hoxc10* and *Hoxc11* were also sequentially activated but their level of expression did not reach that of the genes located at both extremities of the cluster, at least during the period under scrutiny. Therefore, the expression dynamics of *Hoxc* genes in the limb bud ectoderm follows the typical temporal colinear pattern of *Hox* genes expression (32).

In order to validate these results and to determine the spatial distribution of *Hoxc* transcripts in the limb ectoderm, we used *in situ* hybridization (ISH) in tissue sections, which showed a clear restriction of expression to the ectodermal monolayer (Fig. 1C, arrowheads). As the limb bud emerged (E9.5), the expression of 3’-located *Hoxc* genes covered the whole limb bud ectoderm whereas, at later stages (E12.5), the expression of 5’-located genes appeared more restricted to the tip of the limb bud (Fig. 1C). Subsequently, at E14.5, *Hoxc12* and *Hoxc13* transcripts were found concentrated in the digit tips, as detected by ISH both in whole mount and tissue sections and progressively confined to the developing nail region (E16.5) (Fig. 1D-E). In newborn mice, *Hoxc13* expression persisted in the nail matrix while only residual *Hoxc12* transcripts remained in this region (Fig. 1E).

### Function of *Hoxc* genes in nail development

This colinear expression of *Hoxc* genes in the forelimb ectoderm during these developmental stages and the concentration of mRNAs in the nail region suggested a function for these genes in nail morphogenesis. This was previously illustrated both by the mis-shaped nail phenotype displayed by mice lacking the function of *Hoxc13* (12), and by the involvement of *HOXC13* in the human ectodermal dysplasia 9 (ECTD9; OMIM 614931; Lin et al. 2012) characterized by severe nail dystrophy. Consequently, we decided to carefully examine the nail in mice lacking the entire *HoxC* cluster (*HoxC*^−/−^), even though these mutant mice die at birth, presumably due to respiratory problems (14).

Visual inspection of *HoxC*^−/−^ newborn limbs revealed a marked flexion of forelimb digits with a strong suspicion of nail agenesis, based on the lack of the typical light reflection caused by the nail plate, a rich keratinized structure (Fig. 2A). Interestingly, congenital ventral contractures of the digits have also been reported in patients with microdeletions involving the *HOXC* cluster in heterozygosis (34). The complete absence of nails was confirmed by histological analyses. In addition to the lack of nail plate, there was no evidence of the nail matrix, neither of a nail bed and the proximal fold was practically nonexistent, merely reduced to a slight groove (Fig. 2B and 2C, scheme). Therefore, in marked contrast to their wild-type littermates, *HoxC*^−/−^ mutant newborns showed no evidence of nail morphogenesis at the tip of their digits.

**Figure 2.**
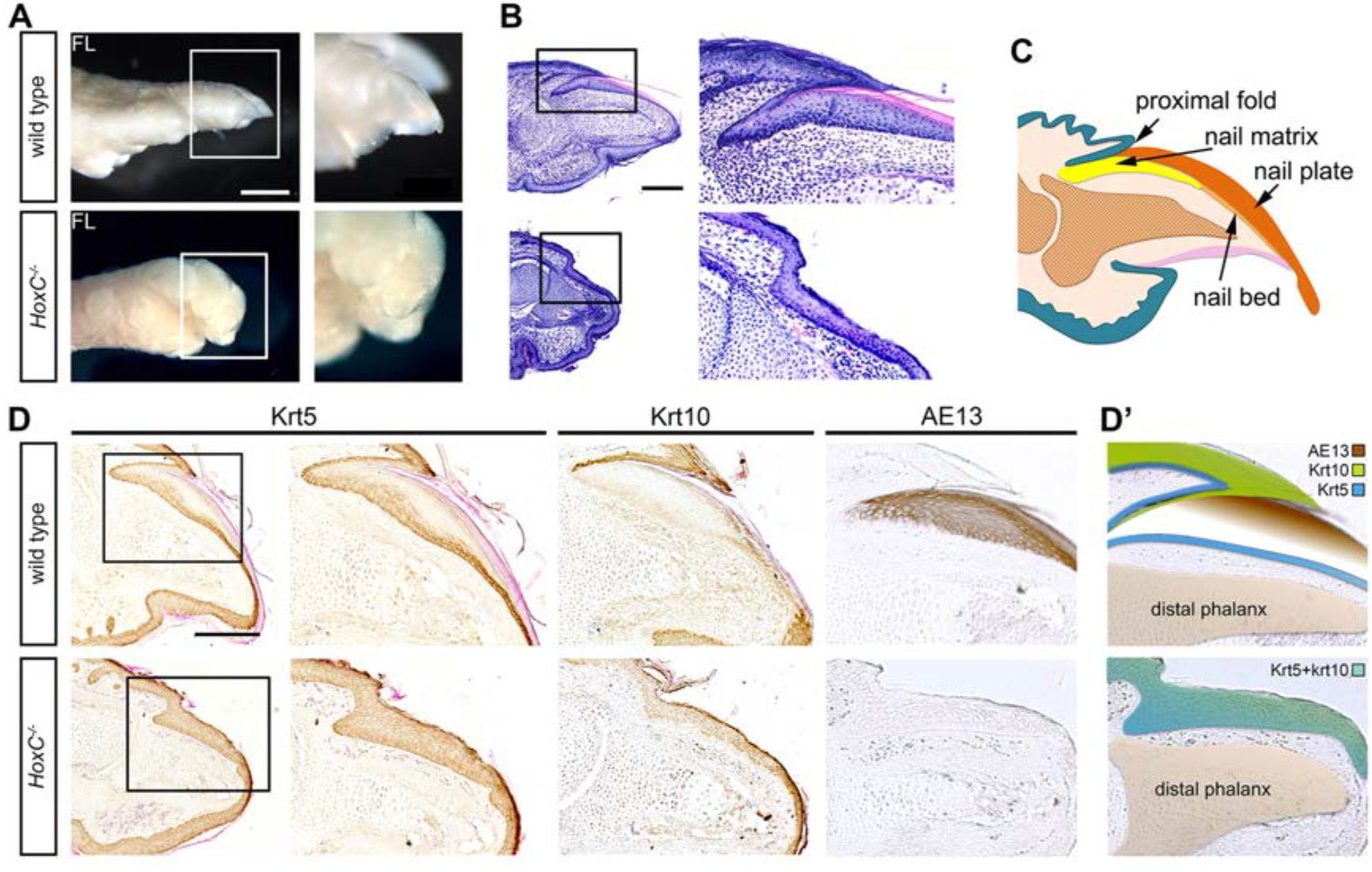
Congenital anonychia in *HoxC*^−/−^ mutant mice. **A)** Photographs showing lateral views of the hand of wild type and *HoxC*^−/−^ newborn mice. Note the pronounced ventral flexion of digits in the mutant specimen. Scale bar 1mm. **B**) Hematoxylin-Eosin stained longitudinal section of a forelimb finger, which shows the absence of the nail organ in *HoxC*^−/−^ mutant animals. Scale bar is 200μm. In A) and B) the framed area is magnified on the right. **C**) Schematic representation of the nail organ showing its various components. **D**) Immunohistochemistry for the detection of Krt5, Krt10 and hard keratins (AE13 antibody) in consecutive longitudinal sections of wild type and *HoxC*^−/−^ newborn mice. For Krt5 immunostaining, a lower magnification is also shown to frame the area under study. Scale bar 200μm. **D’**) Schematic representation of the differentiation state of the nail region in *HoxC*^−/−^ mutants. In all sections dorsal is up and distal to the right.

To evaluate the level of epithelial differentiation in the mutant nail region, we used a range of anti-keratin (Krt) antibodies, which altogether further confirmed the complete absence of the nail organ in *HoxC*^−/−^ mutant specimens. For instance, we observed that Krt5, a specific marker of the basal layer, and Krt10, a supra-basal epithelial marker, unexpectedly persisted in the epithelium of the mutant nail region, while they were normally down-regulated in controls animals (Fig. 2D). In addition, the expression of hard keratins specific for hairs and nails, as detected by using the AE13 antibody, was not found either in the mutant nail region (Fig. 2D). We thus concluded that at birth, the epithelium of the digit tip of *HoxC*^−/−^ mutants displayed a level of differentiation somewhat similar to that of the interfollicular epithelium in control samples, as generally illustrated by the lack of specific nail differentiation (Fig. 2D’, scheme). The analysis of mutant hindlimbs uncovered a similar although slightly milder phenotype (Fig. S2). Again, the expression of hard keratins was not detected while Krt10 abnormally persisted in the digit tip epithelium indicating the impairment of nail differentiation, even though Krt5 was reduced in supra-basal layers. The difference between fore and hindlimb could possibly reflect the expression of additional *Hox* genes in the hindlimb ectoderm, a possibility that remains to be assayed.

This previously unnoticed congenital anonychia observed in *HoxC*^−/−^ mutant animals indicates that *HoxC* cluster genes not only participate to shape the nails, as shown by the *Hoxc13* loss of function mutation (12), but are also necessary for their morphogenesis, at least at the proper time. Indeed, we cannot exclude a developmental delay in the differentiation of the nail organ, for the perinatal death of *HoxC*^−/−^ mutant mice precluded further analysis. In *Hoxc13^−/−^* mice, twisted and fragile nails were reported. At birth, the epithelial differentiation of the nail region, according to Krt5, was similar to normal but the expression of Krt10 persisted in the suprabasal layers of the nail matrix and hard keratins were not detected (Fig. S3). Therefore, it seems that, in the absence of *Hoxc13* function, other *Hoxc* genes can activate the nail differentiation program although the expression of the hard keratins specific to the hair and nails appears to depend on *Hoxc13*.

### Transcriptional regulation of *Hoxc* genes in the ectoderm

The apparent coordinated function of *Hoxc* genes in limb ectodermal derivatives echoed other well described cases where several *Hox* genes located in *cis* are controlled by batteries of common long-acting enhancers positioned outside the respective *Hox* cluster (35–37). We further investigated this possibility by using ATAC-seq (Assay for Transposase-Accessible Chromatin with high throughput sequencing) to identify accessible chromatin regions in isolated ectodermal hulls of E14.5 digit tips (Fig. 3A, first and second tracks). We selected this stage because of the high expression levels of both *Hoxc12* and *Hoxc13*. We looked for open chromatin regions within and around the *HoxC* cluster and identified several open chromatin regions in the *HoxC* cluster itself, which correspond to the activity of *Hoxc* genes, expectedly absent from the control forebrain ATAC-seq dataset. In addition, two focal regions of significant ATAC-seq enrichment were clearly scored outside the *HoxC* cluster in the distal limb ectoderm, whereas they were not detected in the E13.5 forebrain ATAC-seq control dataset (Fig. 3A, third track). Furthermore, these two loci, as well as the *HoxC* cluster, were enriched in H3K4me2 in E10.5 ectoderm, a chromatin mark indicative of transcriptional and enhancer activity (Fig. 3A, bottom track). The presence of other chromatin marks indicative of active enhancer regions using published limb bud datasets was not possible due to the high dilution of ectodermal cells in this material.

**Figure 3.**
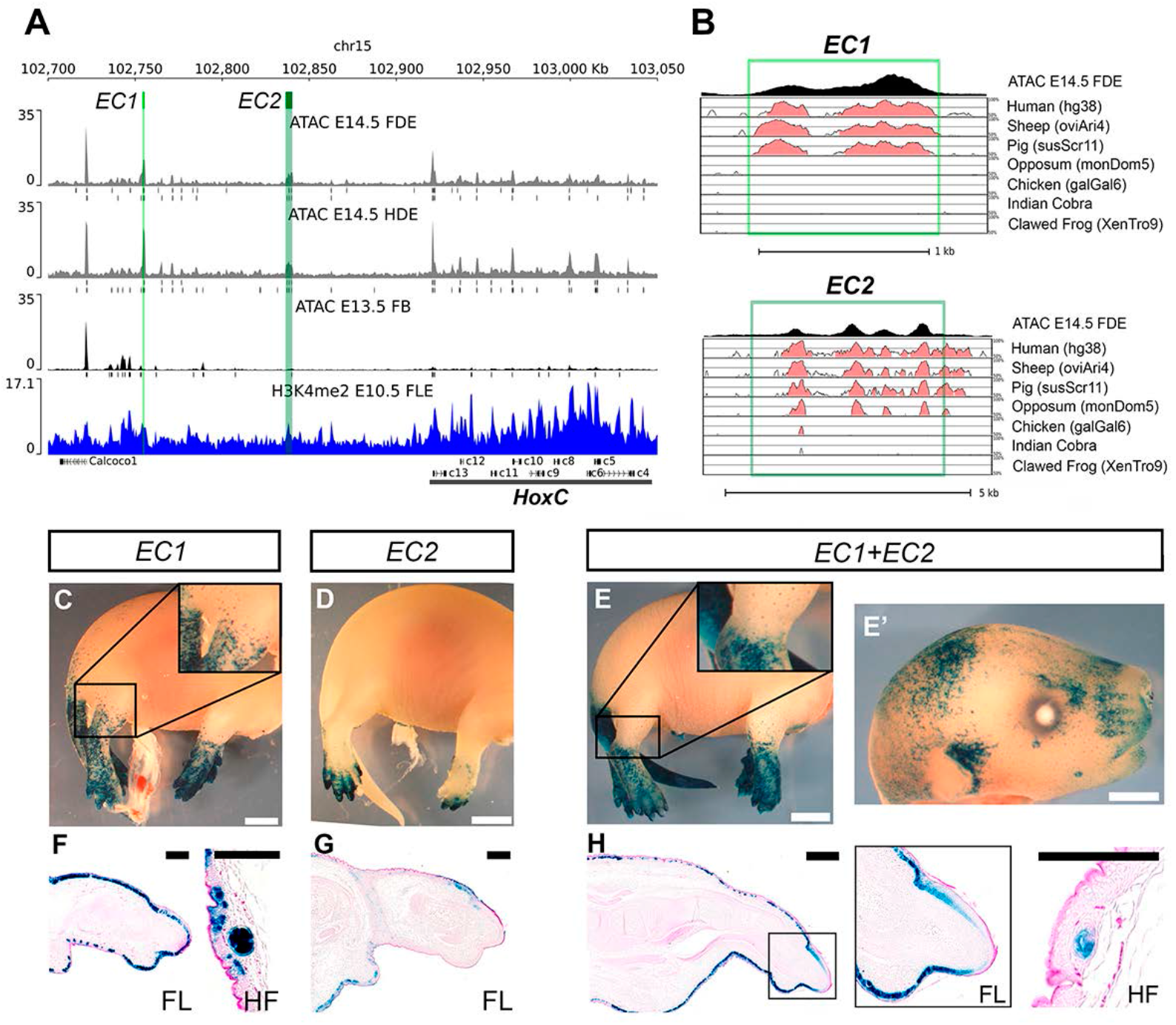
Identification of ectodermal enhancers. **A)** Coverage around the *HoxC* genomic locus of the ATAC-seq of E14.5 forelimb distal ectoderm (FDE) and hindlimb distal ectoderm (HDE) in grey (each one is the mean of two replicates) with the peak calling of each replicate as well as the ATAC-seq of E13.5 forebrain (FB) in black used as a negative control and H3K4me2 ChIP-seq in blue. The two peaks highlighted in green were identified by ATAC-seq only in the distal digit ectoderm but are absent from the control profile in forebrain cells. The ATAC-seq coverage were normalized by million reads in peaks. **B**) Results of the mVISTA tool (shuffle-LAGAN) overlaid with the ATAC-seq coverage track of forelimb distal ectoderm (FDE, top track). Conservation of *EC1* and *EC2* is assessed for 7 different species. For each track the identity over 100bp is shown between 50% and 100%, pink color indicates over 70% of identity. Note that *EC1* is conserved in placental mammals only (light green box), whereas *EC2* is conserved in all mammals (dark green box). We note here the poor quality of the opposum genome for the *HoxC* cluster and the presence of gaps in both cobra and frog genomes. **C-E’**) Reporter activity of *EC1*, *EC2* and *EC1*+*EC2* in E16.5 transgenic fetuses is shown by representative whole-mount *LacZ* expression patterns. Note the prominent *LacZ* staining in the tail and limb bud ectoderm. Scale bars 2mm. **F-H)** Longitudinal sections of limb buds (dorsal up and distal to the right) showing expression restricted to the ectoderm. Magnifications of the hair follicles (HF) in the proximal limb are also shown. Scale bars, 150μm.

These two accessible regions, termed *EC1* and *EC2*, were located 165kb and 81kb upstream of the *HoxC* cluster, respectively, and were highly conserved in their sequences amongst mammals, while absent from other vertebrate species (Fig. 3B). Based on this phylogenetic conservation, the *EC1* and *EC2* sequences were characterized as 1.1kb and 4.1kb large, respectively. Of note, the *EC1* sequence was conserved in placental mammals only, whereas *EC2* was conserved throughout mammals. We set up to assess the activity of these two sequences in transgenic mice either individually or in tandem, using an enhancer *LacZ* reporter construct. Transient transgenic fetuses were harvested at E16.5 and the activity of the reporter *LacZ* gene revealed in whole-mount β-gal staining (Fig. 3C-H, supplementary information).

The *EC1* enhancer displayed a very strong, specific and reproducible activity (5 out of 7 transgenic specimens) in the distal limb bud ectoderm and the developing tail. At the level of the mid zeugopod or the proximal tail, a dense pattern of staining progressively changed to mark hair follicles (HF) exclusively. The HFs were stained over broad areas of the body, although transgene activity was not scored in the whisker pad (Fig. 3C and Fig. S4). The *EC2* sequence elicited a similar activity, yet with a little more variability and smaller penetrance (7 out of 28 transgenic specimens). In general, the ectoderm area showing a dense staining was more restricted to the distal limb and, in most cases, staining was not detected in the tail. While variable areas of head ectoderm were also stained, the transgene did not seem to be active in whiskers, as for *EC1* (Fig. 3D and Fig. S4).

In order to evaluate a potential collaborative or synergistic effect of *EC1* and *EC2*, we assayed their functional activity when introduced in tandem into the enhancer *LacZ* reporter construct. In this combined situation, the *EC1-EC2* transgene displayed very robust and consistent reporter activity in the tail and distal limb ectoderm, as well as in HFs throughout the body surface (10 out of 14 transgenic specimens) (Fig. 3E). Reporter activity was also prominent in patches of head ectoderm, in the whisker pad, and eyelashes (Fig. 3E’). Of note, the dorsal and ventral midlines and the genital tubercle were also stained (Fig. S4).

Longitudinal sections of the limbs of stained embryos for each of the three transgenes confirmed the restriction of their activity to the ectodermal component and illustrated the transition from a distal stronger staining pattern, to a more proximal activity restricted to HFs (Fig. 3F-H). Altogether, these results confirmed that both *EC1* and *EC2* control important aspects of *Hoxc* genes transcription in the ectoderm, both during early embryonic steps and during late phases of differentiation.

### Deletions of the enhancers *in vivo*

To further validate the functional contribution of these two potential enhancer sequences, we used CRISPR–Cas9 genome editing to generate mice lacking each of them, separately (Fig. 4A). Pairs of guide RNAs were used to induce deletions covering either the *EC1* or the *EC2* sequences (Table S3) and zygotes were transformed through electroporation. Stable lines were generated for both the *HoxC*^*delEC1*^ and *HoxC*^*delEC2*^ alleles and bred to homozygosity. Mice homozygous for each deletion were however indistinguishable from their wild-type littermates in all gross aspects such as developmental growth, fertility and viability. Also, their behavior was apparently normal too. Likewise, no significant difference was detected in their ectodermal derivatives and their hair looked healthy and glossy, as well as the nails, which were apparently normal also in the histological analysis (Fig. 4B-C). While more detailed molecular analyses revealed a slightly abnormal ectopic expression of Krt10 in some cells of the nail matrix in *HoxC*^*delEC2*^ homozygous mutants (Fig. 4C), the differentiation of the nail in *HoxC*^*delEC1*^ homozygous mutants appeared completely normal (Fig. 4B). From these results, we concluded that neither *EC1*, nor *EC2* were necessary for the transcription of *HoxC* cluster genes in the ectoderm, at least in the presence of the other enhancer sequence. This raised the possibility that the two sequences carried a redundant specificity and may thus compensate for one another, as was described for a number of related cases in distinct mammalian gene regulatory landscapes (see 35, 38, 39).

**Figure 4.**
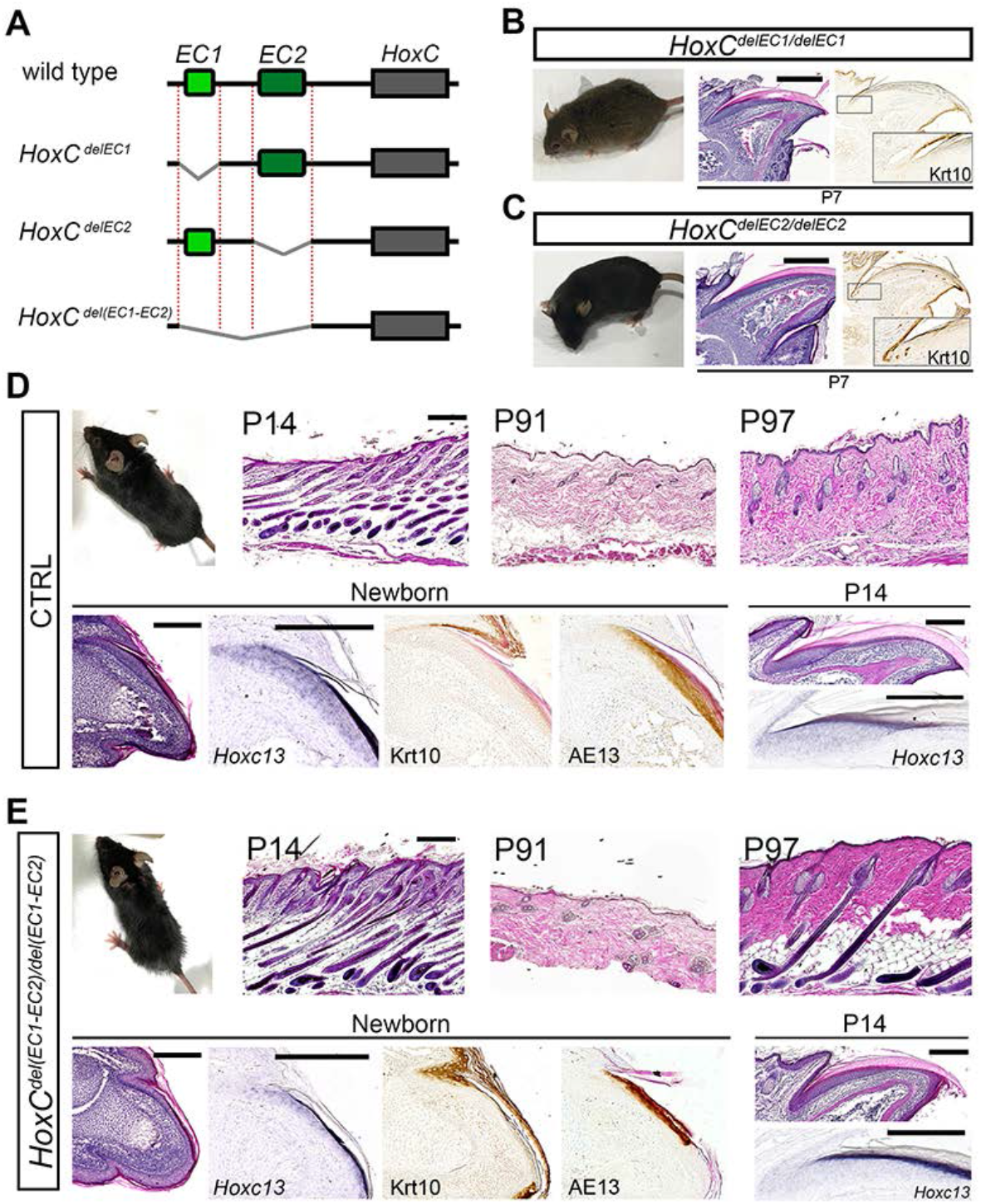
Deletions of enhancers. **A)** Schematic representation of the deletion alleles generated by CRISPR-Cas9. **B)** *HoxC*^*del(EC1)*^ homozygous mice were phenotypically normal and showed wild type nail development **C)** *HoxC*^*del(EC2)*^ homozygous mice appeared normal as well, even though a minor ectopic expression of Krt10 was detected in the nail matrix. Scale bars in A and B, 400μm. **D-E)** Mice homozygous for the deletion of both enhancers (*HoxC*^*del(EC1-EC2)/del(EC1-EC2)*^) displayed disheveled fur and hypoplastic nails. Histological comparison of hair follicles (HFs) in HE-stained sections of dorsal skin from P14 control and *HoxC*^*(delEC1)*^ homozygous mice showed some hair shafts distorted at the level of the sebaceous glands. Over time, however, an acceleration in the cycling rate became evident as illustrated by mutant dorsal HFs having precociously entered the next anagen at P97 while control HFs remain in telogen as in P91. Scale bars in D and E, 200μm. At birth, the mutant nails are hypoplastic as indicated by the abnormal expression of Krt10 remaining in the proximal nail matrix and the substantial reduction in *Hoxc13* and hard keratins expression. The nail defect was highly attenuated over the first two weeks of postnatal development (P14) despite continued reduction in *Hoxc13* mRNA levels.

To investigate whether *EC1* and *EC2* may indeed share parts or all of their spatio-temporal regulatory specificities, we deleted a 86kb large DNA sequence containing the two sequences to produce the *HoxC*^*del(EC1-EC2)*^ allele (Fig. 4A). Mice homozygous for this allele were viable and fertile. While they globally looked healthy, they exhibited mild but consistent fur and nail phenotypes (Fig. 4D-E). When compared to control littermates, homozygous *HoxC*^*del(EC1-EC2)*^ mice showed the persistence of a disheveled fur (a rough fur phenotype) with some of the mice developing bald patches on their backs. Histological analyses of the skin did not reveal any obvious defect in the morphogenesis of the HFs but showed some hair shafts distorted at the level of the sebaceous glands, reminiscent but less marked than those observed in *Hoxc13*-null and *Foxn1*-null mice and that associate with defective keratinization of the hair shaft (40). Over time, after the second or third hair cycles, an acceleration in the hair cycle became obvious. Therefore, when compared to wild type littermates, homozygous *HoxC*^*del(EC1-EC2)*^ mutants showed a precocious re-initiation of the next cycle with shortened telogen and premature anagen development, as shown in the analysis of P91 and P97 littermates (Fig. 4D-E). This is possibly driven by the loss of hair inducing a plucking effect similar to the situation described in *Foxn1*-null mice (41). The nails were unusually hypoplastic at birth, showing abnormal Krt10 expression in the proximal nail matrix as well as a substantial reduction in hard keratins expression (Fig. 4D-E). Accordingly, *HoxC*^*del(EC1-EC2)/del(EC1-EC2)*^ mutant specimens displayed a substantial reduction in the amount of *Hoxc13* mRNAs (Fig. 4D-E). The nail defects became attenuated during the first two weeks of postnatal life and, by P14, the histology of the nail was close to normal. These observations suggested that the presence of both *EC1* and *EC2* is required to reach a threshold in *Hoxc* genes mRNAs that is needed for the normal morphogenesis of epidermal organs.

To further document a possible role of *Hoxc* gene dosage in the development of ectodermal organs and to examine the function of *EC1* and *EC2* in this context, we looked at the phenotypic consequence of deleting the two *EC1* and *EC2* enhancers in a sensitized genetic background, where the expression of the presumptive target *Hoxc* genes was further reduced. We generated trans-heterozygous mice harboring either one of the single or multiple enhancer deletions, over the full deletion of the entire *HoxC* cluster. Animals of the *HoxC*^*del(EC1)*/−^ or *HoxC*^*del(EC2)*/−^ genotypes showed no particular phenotype, indicating that the individual functional contribution of either the *EC1* or the *EC2* enhancers was dispensable even under conditions of sensitized background.

However, the removal of both enhancers on this same background (*HoxC*^*del(EC1-EC2)*/−^) led to a striking phenotype characterized by the total absence of body hair (Fig. 5A). Trans-heterozygous *HoxC*^*del(EC1-EC2)*/−^ mice were born at the expected mendelian ratio, were viable and fertile and displayed a normal behavior. The phenotype became readily detectable shortly after birth due to their nude aspect (Fig. 5A). In contrast to heterozygous or wildtype control mice, no fur coat emerged from the skin of *HoxC*^*del(EC1-EC2)*/−^ specimens during the first postnatal week, reminiscent of the phenotype of *Hoxc13* homozygous mutant animals (12). Interestingly, despite their lack of hairs, they displayed vibrissae at birth, in contrast to *Hoxc13* mutant mice. These mice remained nude until the new anagen phase, at about five weeks of age, when they developed hairs in a cranio-caudal wave. The hairs were rapidly lost in the next couple of days, however, and the mice remained nude until the next hair cycle when a new wave of hairs emerged and was rapidly lost again. Thereafter, the waves of hair regrowth and loss became more irregular. This iterative pattern of hair formation and loss resulted in a typical cyclic alopecia phenotype (Fig. 5A).

**Figure 5.**
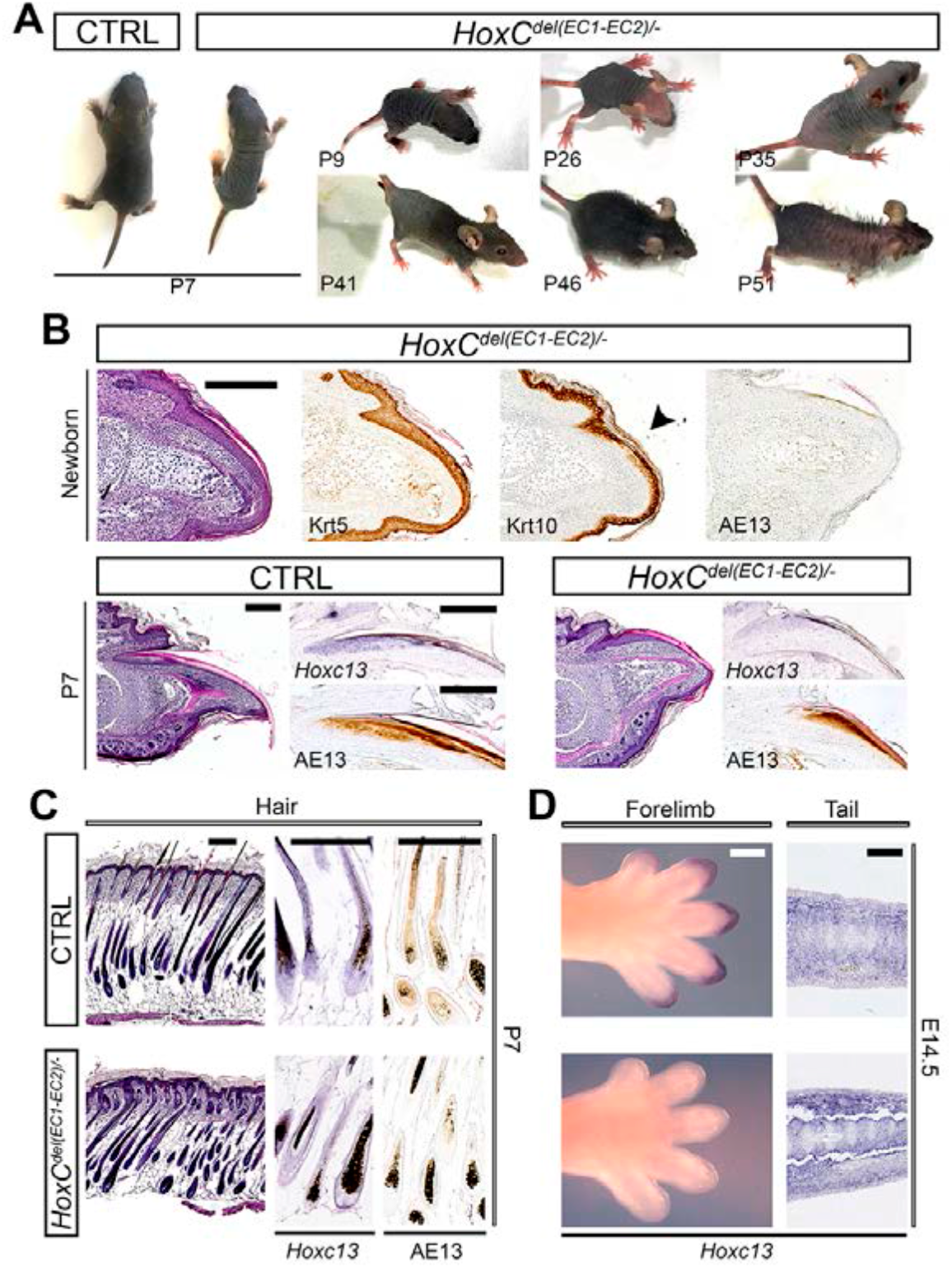
Detectable effects of *EC1* and *EC2* combined deletion in a sensitized background. **A)** *HoxC*^*del(EC1-EC2)*/-)^ trans-heterozygous mice lack external hair during the first hair cycle and develop cyclic alopecia. **B)** When compared to control littermates (*HoxC*^*del(EC1-EC2)*/+^), the nail was found under-developed in such mutant mice, with only minor focal areas of specific nail differentiation at birth (arrowhead). The phenotype progressively attenuated with age, as nail differentiation in the distal nail matrix, as revealed by the expression of hard keratins, was observed in P7 mutants, but always accompanied by a reduced level of *Hoxc13* mRNAs. Scale bars 200μm. **C)** Histological analysis of HFs of dorsal skin from P7 mutant mice showed a number of HFs that became distorted when entering the epidermis. This was accompanied by a marked reduction of hard keratins detected by AE13 immunohistochemistry, and by a downregulation of *Hoxc13* expression. **D)** At E14.5, *Hoxc13* expression was bellow detection level in the ectoderm of the digit tips, yet it normally persisted in the tail mesenchyme. Scale bars 400μm.

At birth, the nails of *HoxC*^*del(EC1-EC2)*/−^ mice were severely underdeveloped with only minor focal areas of specific nail differentiation and a total absence of nail plate and hard keratin expression (Fig. 5B, compare with control in Fig. 2D and 4D), reminiscent of the *HoxC*^−/−^ nail phenotype (Fig. 2). Over time (P7), however, the defect became less pronounced although the expression of *Hoxc13* and hard keratins remained lower than normal. The nail and hair defects observed in *HoxC*^*del(EC1-EC2)*/−^ mice are strikingly similar to that reported for mice lacking the *Foxn1* function, a gene known to be a transcriptional target of *Hoxc13* and in turn, a transcriptional regulator of hard keratins of the hair shaft (40, 42). Accordingly, the histological analysis of hair follicles showed a phenotype similar to that described in *Foxn1*-null mice. Hair follicles formed normally during fetal development but the hair shaft was distorted when entering the epidermis (Fig. 5C), a phenotype that was considered to derive from the lack of hard keratins in *Foxn1* mutants. Hard keratins were reduced substantially in the HFs of *HoxC*^*del(EC1-EC2)*/−^ mice, correlating with a lower level of *Hoxc13* mRNAs.

The CRISPR–Cas9 approach also triggered an inversion of the targeted region producing the *HoxC*^*inv(EC1-EC2)*^ allele. Mice homozygous for this allele (*HoxC*^*inv(EC1-EC2/inv(EC1-EC2)*^) were indistinguishable from wild type littermates and displayed normal fur and nails indicating that the inverted rearrangement of the enhancers had no functional consequence (Fig. S5). In addition, mice trans-heterozygous for this inverted configuration and the deletion of the *HoxC* cluster (*HoxC*^*inv(EC1-EC2)*/−^) did not show any abnormality (Fig. S5).

Altogether, our results suggest that both *EC1* and *EC2* contribute to the normal level of transcription of *Hoxc13* in the ectoderm, likely in association with other, as yet uncharacterized enhancers. In contrast to both *Hoxc13*^−/−^ (12, 13) and *HoxC*^−/−^ (14) mutant mice, the hereby described *HoxC*^*del(EC1-EC2)*/−^ mice were viable and healthy, indicating that the induced defects were likely localized to the reported ectodermal components. We further tested this hypothesis by looking at the expression of *Hoxc13* at E14.5, both in the limb ectoderm and in the growing tail mesenchyme, another site where *Hoxc13* is normally transcribed. While no detectable *Hoxc13* transcripts were scored in the limb ectoderm by whole-mount *in situ* hybridization, expression of this gene in the tail mesenchyme remained as in controls (Fig. 5D), supporting the definition of *EC1* and *EC2* as enhancers carrying a predominant ectodermal specificity.

## Discussion

In this work, we report that the *HoxC* cluster has evolved a general functional specificity for some ectodermal organs, the nails and hairs, and that *Hoxc* genes are activated in the embryonic limb ectoderm following a colinear time sequence. We also show that part of this specificity is achieved through the action of two enhancers positioned at a distance from the gene cluster itself and acting in combination to raise the transcription level of several *Hoxc* genes in *cis*.

### Colinear and non-colinear expression of *HoxC* cluster genes in the ectoderm

During the formation and elongation of the major body axis, *Hox* genes belonging to all four gene clusters are activated in a time sequence that reflects their topological order along their respective clusters (32). While the impact of this temporal colinearity upon the colinear distribution of *Hox* transcripts in space is still a matter of discussion (see e.g. 43–46), the mechanism underlying this phenomenon is accompanied by a progressive transition from a negative to a transcriptionally permissive chromatin structure (47). This phenomenon is however observed only during the first wave of activation of a given *Hox* cluster. Indeed, once the cluster open, global enhancers located in the surrounding regulatory landscapes can subsequently regulate sub-groups of genes located in *cis* simultaneously, as was shown for example for the transcription of *Hoxd* genes in the developing digits or external genitals (48).

Here we show a clear collinear time-sequence in the activation of *Hoxc* genes in the limb ectodermal component. This temporal sequence in mRNAs production does not correspond to any distinct ectodermal structure or compartment, which could have required the proper deployment of various HOXC proteins in time and space, for example to identify different cell types. We interpret this as an indication that the *HoxC* cluster was initially not transcribed at all in the ectodermal precursors and hence that its chromatin structure was not transcriptionally challenged before this precise spatio-temporal situation. As a consequence, upon activation, the gene cluster had to be processed as all *Hox* clusters during their initial phase of activation, following a 3’ (*Hoxc4* in this case) to 5’ (*Hoxc13*) direction. In this view, the temporal colinear activation we clearly detect in the limb ectoderm may not reflect any particular functional constraint other than allowing eventually the transcription of the most 5’ located genes that seem to be required for nail development. As such, it may illustrate an intrinsic constraint of the system itself (49). Alternatively, temporal colinearity in limb ectoderm may underly the slightly different distributions of *Hoxc* genes, which in turn may impact the morphology of the nails.

In contrast, expression of *Hoxc* genes in hair follicles occurs throughout the body, with no particular rostral to caudal colinear distribution. For this reason, it was initially proposed to ‘break’ spatial colinearity (12). In fact, this deviation from the spatial colinearity observed during the development of the major body axis is only detected in the ectodermal compartment, since *Hoxc* gene expression in the dermal papilla seems to be colinear indeed, with *Hoxc4* to *Hoxc10* transcribed in dorsal skin, whereas *Hoxc10* to *Hoxc13* mRNAs are found in the tail (19). Altogether, our results further document the functional co-option of at least *Hoxc13* (see below) for a late but critical function in the hair and nail epidermal organs. Of note, the two enhancers reported here are located outside of the *HoxC* cluster, a global regulatory strategy often associated to *Hox* clusters and observed in many other developmental contexts (see 37, 50–52).

### Sub-functionalization of *Hox* clusters

All four *Hox* gene clusters in amniotes are involved in the specification of structures during the development of the major body axis, which is likely their most ancestral function. In addition to this function, conserved in most animals with a bilateral symmetry, clusters were subsequently co-opted for specific functions, after the rounds of genomic duplications (53, 54). For example, both the *HoxD* and *HoxA* clusters are necessary to properly develop limbs and external genitalia. Such cluster-specific functions may be redundant between gene clusters (as is the case in limbs and genitals) and hence the full assessment of such global specificities has been difficult to evaluate due to the experimental problems to produce multiple full *Hox* cluster deletions (see 8).

In the case of *HoxC*, however, its deletion alone induced a drastic phenotype associated with the ectodermal compartment, thus suggesting that no other *Hox* cluster is equally functional in these important organs and could then have compensated for the loss of these genes. This is re-enforced by the fact that only *Hoxc* genes were found clearly expressed in the developing limb ectoderm (Figure 1A). This observation suggests that the functional co-option of the *HoxC* cluster into ectodermal organs occurred after the full set of genome duplications, potentially to take over the control of structural components of ectodermal tissue such as the hard keratins.

Without an extensive genetic analysis, it is difficult to determine how many *Hoxc* genes, and which ones, were involved in these two functional recruitments. Also, the situation may be slightly different for the nails and the hair follicles, indicating either distinct functional requirements for these two organs, or/and different modalities in the recruitment and implementation of enhancers. In the case of hair follicles, indeed, *Hoxc13* function seems to be sufficient as *Hoxc13*-null mice are nude (12, 13) and the analysis of these mutant mice revealed the function of this gene in the production of hard keratins and keratin-associated-proteins that are direct targets of *Hoxc13* (40, 55). In the absence of hard keratins, the hair shafts are brittle and readily break as they emerge through the skin surface, thus producing the nude phenotype. Likewise, the nails are fragile and twisted (40, 56). The *Foxn1* gene, whose null mutation generates a nude phenotype, was also described as a *Hoxc13* regulatory target during hair and nail differentiation (40, 42, 57).

In the case of the nails, the functional contribution of *Hoxc* genes seems to be slightly different than in the hair. Indeed, while nails of *Hoxc13* mutant mice are fragile (12, 13), we show here that their abnormal phenotype is more severe in the complete absence of the *HoxC* cluster. *HoxC*^−/−^ mutant animals, which die as newborns due to defective lung development (14), show a congenital anonychia with no trace of nail specific keratinocyte differentiation, a phenotype that remained unnoticed. Therefore, other *Hoxc* genes, possibly *Hoxc12*, whose expression is similar to that of *Hoxc13* (18), could either have a function during nail development, or partially substitute for *Hoxc13* function when the latter one is abrogated.

### A mammalian-specific (enhanced) regulation?

The critical importance of *Hoxc13* in mammalian hairs and nails is also emphasized by the strikingly similar phenotypes resulting from loss of function conditions in different species including rabbits and pigs (33, 58, 59). In these cases, hair and nail differentiation is impaired in a way comparable to the murine situation. In addition, there are clear similarities with the human ectodermal dysplasia 9 (ECTD9, OMIM #614931), also caused by the loss of function of *HOXC13*. Ectodermal dysplasias (ECTDs) are a group of congenital disorders characterized by abnormal development of two or more ectodermal appendages, without other obvious systemic anomalies. Hairs are most commonly affected in association with alterations of the nails, teeth (anodontia or hypodontia) or sweat glands. Amongst the very rare ECTDs involving hairs and nails exclusively is the ECTD9, which is caused by mutations in *HOXC13* (33, 60, 61).

While hairs are specific for mammals, different skin appendages exist in other vertebrates, which also express *Hox* genes during their development, like in feather buds (62). Terminal keratinized digit structures like nails also exist in all tetrapods, under a variety of forms and, in the chick, expression of *Hoxc13* was observed in the tips of digits along with the development of claws (Fig. S6). Therefore, it is likely that a sustained level of *Hoxc* genes is required throughout tetrapods, in conjunction with several ectodermal organs that involve high amounts of keratins, regardless of the final morphologies of these organs (hairs or feathers, nails or claws). Accordingly, *Hoxc* genes could be necessary to activate a ‘keratinization program’, which would be interpreted differently depending on the phylogenetic context. The two global enhancer sequences we report in this study may be necessary for this ectodermal specific-activation of *Hoxc* genes, as suggested by their behavior when introduced into transgenic mice.

This view is nevertheless difficult to reconcile with the fact that neither of these two ectodermal enhancers seems to be conserved outside mammals. For instance, it is not found in birds, neither at the syntenic position nor elsewhere, while one would expect the same ectodermal regulation to occur for this highly conserved gene cluster. One possible explanation to this paradox may rely upon the quantitative *versus* qualitative aspect of this regulation. Indeed, it is clear from our results that the removal of these two sequences together did not entirely abolish *Hoxc* gene expression in the concerned ectodermal tissues. In this case, hypomorph phenotype was observed in newborns, which in the nails was progressively corrected during the next two weeks. This strongly indicates that the dose of HOXC13 protein is directly related to the final morphology of the nails and that a quantitative deficit in protein can be slowly corrected by reaching the required amount over time.

To identify the limb bud cell subpopulations with *EC1* and *EC2* chromatin accessibility, we quantified their accessibility into the single cell ATAC-seq (scATAC-seq) dataset recently generated from E11.5 mouse forelimb buds (63). The representation of this quantification on the previously calculated uniform manifold approximation and projection (UMAP) coordinates showed that most of the cells having these enhancers open were ectodermal cells (Fig. S7). In addition, all those cells but one showing ATAC-seq signals on both enhancers were ectodermal cells. These results thus support the conclusion that both *EC1* and *EC2* are specific ectodermal enhancers and that they can function in the same cell, suggesting a collaborative or synergistic mechanism.

This quantitative aspect was also demonstrated by our genetic analyses using other alleles of *HoxC*, which revealed that the *EC1* and *EC2* enhancers control only part of the steady-state level of *Hoxc* mRNAs in ectodermal organs, thus indicating that the ectodermal expression of *Hoxc13* (at least) is partly controlled by other sequences located outside the *HoxC* cluster, which would also contribute to the *Hoxc13* expression level. In this context, it is possible that the requirement for HOXC proteins to accompany the development of ectodermal organs be slightly different in various classes of vertebrata, with a higher protein level necessary in mammals than in birds. It is thus conceivable that the *EC1* and *EC2* enhancers evolved along with the mammalian lineage to progressively adjust the overall dose of HOXC proteins such as to achieve well adapted ectodermal organs. In this scenario, these enhancers would thus be specific for mammals, but involved into a function much more largely distributed within vertebrates.

## Materials and Methods

A more detailed description is provided in Supplementary methods

### Mouse strains and animal ethics

We have used the four *HoxC*^*del(EC1)*^, *HoxC*^*del(EC2)*,^ *HoxC*^*del(EC1-EC2)*^ and *HoxC*^*inv(EC1-EC2)*^ mutant lines generated here by CRISPR/Cas9 and the previously published *Hoxc13*^*GFP*^(64) and *HoxC*-null (14) strains. All animal procedures were conducted according to the EU regulations and the 3R principles and were performed in agreement with the Swiss law on animal protection (LPA), under license No GE 81/14 (to DD) and reviewed and approved by the Bioethics Committee of the University of Cantabria.

### Histological analysis, immunohistochemistry and *in situ* hybridization

Dissected tissues were fixed in 4% PFA and processed for paraffin embedding and microtomy. Hematoxylin-Eosin staining, immunohistochemistry and *in situ* hybridization were performed following standard procedures. The primary antibodies used were: anti-Keratin 5 (SIGMA, SAB4501651), anti-Keratin 10 (Biolegend, 905401) and anti-Pan-Cytokeratin (AE13; Santa Cruz, sc-57012).

### RNA-seq

For each of the 16 samples, RNA libraries were prepared using the Truseq^®^ Stranded mRNA Library Prep Library Preparation kit (Illumina, 20020594) and 100-bp single reads were generated.

### ATAC-seq and mouse transgenic enhancer assays

We performed ATAC-seq (66) on isolated ectodermal cells from the distal tips of wild-type E14.5 embryos. Two forelimb and two hindlimb biological replicates were generated. ATAC-seq on E13.5 forebrain was used as control.

The analysis of the conservation of the *EC1* and *EC2* enhancers was performed with mVISTA shuffle-LAGAN (67). The activity of *EC1, EC2* and *EC1*+*EC2* was assayed in mouse transgenesis by using a vector carrying a β-globin minimal promoter and the *LacZ* coding sequence (pSK-LacZ).

### CRISPR-Cas9 modifications

The gRNAs used to generate the *delEC1, delEC2, del(EC1-EC2)* and *inv(EC1-EC2)* mutant lines were designed with CHOPCHOP (https://chopchop.cbu.uib.no/). The gRNA sequences used in the CRISPR experiments are listed in Table S3. The CRISPR-Cas9 methodology used to generate *delEC1* and *delEC2* mouse alleles was adapted from Qin *et al* (68) and the *del(EC1-EC2)* and *inv(EC1-EC2)* mouse alleles were generated with the Alt-R^®^ CRISPR-Cas9 System from IDT (https://eu.idtdna.com/pages/products/crispr-genome-editing/alt-r-crispr-cas9-system). The mutant lines were generated by electroporation. See SI for more information.

B6CBAF1/J Fertilized eggs were collected from the oviducts of E0.5 pregnant females. The collected eggs cultured in WM medium were washed with Opti-MEM (Gibco, 31985-047) three times to remove the serum-containing medium. The eggs were then lined up in the electrode gap filled with the electroporation solution, electroporated and transferred into pseudo-pregnant foster mice (68, 69).

## Data availability

All raw and processed datasets of RNA-seq, ATAC-seq are available in the Gene Expression Omnibus (GEO) repository under accession number GSE150702. All sequences used for conservation analysis as well as all bioinformatics scripts needed to reproduce the figures from raw data are available at https://github.com/lldelisle/scriptsForFernandezGuerreroEtAl2020. The raw and process datasets of the H3K4me2 ChIP-seq is registered under GSM4294458.

## Acknowledgments

We thank Sara Lucas, Bea Romero, Mar Rodriguez and Bénédicte Mascrez for their help with electroporation of embryos and handling crosses as well as Laura Galán for excellent technical assistance. This work was supported by funds from the Ecole Polytechnique Fédérale (EPFL, Lausanne), the University of Geneva and the Swiss National Research Fund (No. 310030B_138662) and the European Research Council grants Regul*Hox* (No 588029) to DD and by the Spanish Ministry of Science and Innovation (Grant BFU2017-88265-P) to MAR.

## Supplementary Information

### Supplementary Materials and Methods

#### Mouse strains

The *Hoxc13*^*GFP*^ (1) and *HoxC*-null (2), and the newly produced *HoxC*^*delEC)*^, *HoxC*^*delEC2*,^ *HoxC*^*del(EC1-EC2)*^ and *HoxC*^*inv(EC1-EC2)*^ mutant lines were used in this study. All strains were maintained as backcrosses with a B6XCBA mixed background. Genotyping was performed using tail biopsies or embryonic membranes. Embryos of the desired embryonic day were obtained by cesarean section. The primers used for genotyping the new generated alleles are listed in Table S1.

#### Hematoxylin-Eosin staining

Tissues were fixed in 4% PFA, dehydrated and embedded in paraffin following standard procedures. Seven microns tissue sections were rehydrated and stained with Harris Hematoxylin (Merck, HX57998853) and Eosin Y (SIGMA, E4382-25G). After staining, sections were dehydrated in graded ethanol and mounted in cytoseal60 (Richard-Allan Scientific, 8310).

#### Whole-mount and section *in situ* hybridization and immunohistochemistry

*In situ* hybridization was performed in whole mount or paraffin sections with digoxigenin-labeled antisense riboprobes following standard procedures. The probes used were mouse *Hoxc5*, *Hoxc8*, *Hoxc12* and *Hoxc13* and chick *Bambi* and *Hoxc13* (3, 4)(Table S2). Immunohistochemistry was performed in paraffin sections (7 μm) after antigen retrieval with citrate buffer 10mM pH=6.

#### Transgenic mice for LacZ activity

The genomic region containing the *EC1* enhancer (mm10, chr15:102,754,343-102,755,452) and *EC2* (mm10, chr15:102,836,392-102,840,307) were PCR-amplified using specific primers containing restriction sites for subsequent cloning into the pSK-LacZ vector 5’ to the β-globin minimal promoter. The insert carrying the putative enhancer sequence, β-globin minimal promoter and LacZ coding sequence was excised from the vector backbone by digestion with ApaI–XbaI.

The fragment was gel-purified and injected into the pronucleus of fertilized oocytes that were transferred to foster CD1 females. Transient transgenic embryos were harvested at E16.5 and processed for detection of β-galactosidase activity.

#### β-Gal staining

Embryos were dissected in cold PBS and yolk sac samples were collected for genotyping. Embryos were fixed 20 min at RT in Fix Solution (1x PBS pH=6.8, 2mM MgCl2, 4% PFA, 0.2% Glutaraldehyde (SIGMA, G5882), 5mM EDTA). After rinsing in Wash solution (1x PBS pH=6.8, 2mM MgCl2, 0.2% NP40, 0.01% sodium deoxycolate (DOC)), the embryos were washed 5×20min in Wash Solution. LacZ signal was revealed with staining solution (5mM K3[Fe(CN)6], 5mM K4[Fe(CN)6]), 0.5 mg/mL X-gal (Thermo, R0402)/DMF(SIGMA, D4551) at 37°C overnight. After staining samples were washed once in wash solution and fixed again in 4% PFA/PBS at 4°C overnight. To analyze the precise localization of lacZ signal in the limbs, stained limbs were processed for paraffin embed, sectioned and lightly stained in Eosin.

#### Dissection of samples for RNA-seq and ATAC-seq

For the RNA-seq, the distal stripe (150 microns width) of E9.5, E10.5, E11.5 and E12.5 forelimb buds was dissected out in cold PBS and incubated in 0.25% trypsin (SV30037.01, GE Healthcare, Logan, Utah) on ice for 20 min. The distal limb stripes were then transferred to cold PBS where the ectoderm was separated from the mesoderm with fine forceps. Two biological replicates were collected for each stage and tissue and the same batch of embryos was used for the ectoderm and mesoderm corresponding replicates. From each sample, total RNAs were purified using the RNeasy Plus Mini Kit (QIAGEN, 74134) for the first replicates and RNeasy Mini Kit (QIAGEN, 74104) for the second replicates.

For the ATAC-seq, two forelimb and two hindlimb biological replicates were generated, each replicate consisting of the ectoderm of E14.5 digits tips from wild type embryos. The digit tips were dissected and the ectoderms isolated from the subjacent mesoderm by trypsin digestion (0.25%, 40 min., on ice) and then dissociated to single cell suspension by pipetting up and down. E13.5 forebrain samples were collected as a control. Forebrain portion was removed from the skull cartilage with forceps and digested in PBS with 10% FCS and digested with collagenase at 37°C at 400rpm for 20 minutes.

#### RNA-seq analysis

The gtf from Ensembl version 93 was filtered to remove read through transcripts and all non-coding transcripts from a protein-coding gene. In addition, all genes with the same gene name which overlaps were merged under the same gene to avoid ambiguous reads. Adapters and bad-quality bases were removed with Cutadapt version 1.16 options -m 15 -a GATCGGAAGAGCACACGTCTGAACTCCAGTCAC -q 30. Reads were further mapped using STAR version 2.6.0c with ENCODE parameters with the gtf file described above. FPKM values were computed with cufflinks version 2.2.1 omitting mitochondrial genes. Strand specific coverage was computed with bedtools version 2.27.1 from uniquely-mapped reads and normalized to the number of million uniquely mapped reads for each RNA-seq dataset. For Hierarchical Clustering Analysis (HCA) and Principal Component Analysis (PCA) only the 500 genes with the higher variance were considered, using log2(1 + FPKM) values. The correlation matrix was performed using Euclidean distance with ward clustering. Figure 1A, S1B-D were done with R (www.r-project.org). Figure 1B was done with pyGenomeTracks (5).

#### ATAC-seq

Adapters and bad-quality bases were removed with Cutadapt version 1.16 options -m 15 -a CTGTCTCTTATACACATCTCCGAGCCCACGAGAC -A CTGTCTCTTATACACATCTGACGCTGCCGACGA -q 30. Reads were further mapped using bowtie2 version 2.3.5 options --very-sensitive --no-mixed –no-discordant --dovetail -X 1000. Only pairs with MAPQ30, concordant and in chromosomes which are not chrM were kept. Duplicates were removed with picard and the bam file was converted to bed with bedtools. Peak calling and coverage were obtained with macs2 version 2.1.1.20160309 with options --nomodel --keep-dup all --shift -100 --extsize 200. For each dataset, summit positions were extended 500bp each side and merged and the number of reads falling into these regions was used to normalize the coverage. Figure 3A was done with pyGenomeTracks (5).

#### Conservation of *EC1* and *EC2*

For the 7 species and genome versions: human (hg38), sheep (oviAri4), pig (susScr11), oppossum (monDom5), chicken (galGal6), Indian Cobra (6) and the clawed frog (XenTro9), the fasta containing the sequence between *Hoxc13* and *Calcoco1* was extracted (except in galGal6 were there was no *Calcoco1*, thus the region was extended to *Atp5g2* and in monDom5 where only *Hoxc5* and *Hoxc6* were annotated thus 400kb starting at *Hoxc5* were used). All these sequences with the two regions centered on *EC1* and *EC2* 1.5 times their size were aligned using mVISTA shuffle-LAGAN (7).

#### CRISPR-Cas9 modifications

The gRNAs used to generate the *delEC1, delEC2, del(EC1-EC2)* and *inv(EC1-EC2)* mutant lines were designed with CHOPCHOP (https://chopchop.cbu.uib.no/). The target regions and gRNA sequences used in the CRISPR experiments are listed in Table S3.

The CRISPR-Cas9 methodology used to generate the *delEC1* and *delEC2* mouse lines was adapted from Qin *et al*. (8): The px330 plasmid carrying the wild-type (WT) Cas9 (9) was used as the DNA template for amplification of the Cas9 coding sequence in a polymerase chain reaction (PCR). The T7 promoter sequence was added to the forward primer and reverse primer from the coding sequence of the Cas9 gene. PCR product was amplified using the AccuPrime PCR system (Life Technologies, 12339016) and purified using the QIAquick PCR purification kit (Qiagen, 28104) and *in vitro* transcription (IVT) performed using the mMESSAGE mMACHINE T7 ULTRA Transcription kit (Life Technologies, AM1345). For sgRNA synthesis, the T7 promoter sequence was added to sgRNA template/forward primer and the IVT template generated by PCR amplification. The T7-sgRNA PCR product was purified and used as the template for IVT using MEGAshortscript T7kit (Life Technologies, AM1354). Both the Cas9 mRNA and the sgRNAs were purified using the MEGAclear kit (Life Technologies, AM1908). Aliquots from an IVT reaction were separated on agarose gel to assess quality from a reaction. Single-stranded oligos were ordered as PAGE Ultramer from Integrated DNA Technologies.

The most 5’ crRNA used to generate *delEC1* allele and the most 3’ crRNA used to generate the *delEC2* allele were used jointly to generate both the *del(EC1-EC2)* and *inv(EC1-EC2)* mouse alleles. Following the IDT protocol for the Alt-R^®^ CRISPR-Cas9 System, crRNA and tracrRNA of interest were resuspended in Nuclease-Free Duplex Buffer and mixed to form the crRNA:tracrRNA duplex. The crRNA:tracrRNA duplex was combined with the Alt-R Cas9 Nuclease 3NLS (IDT, 1074181) in Opti-MEM medium to form the ribonucleoprotein (RNP) complex.

## Supplementary Figures

**Figure S1.**
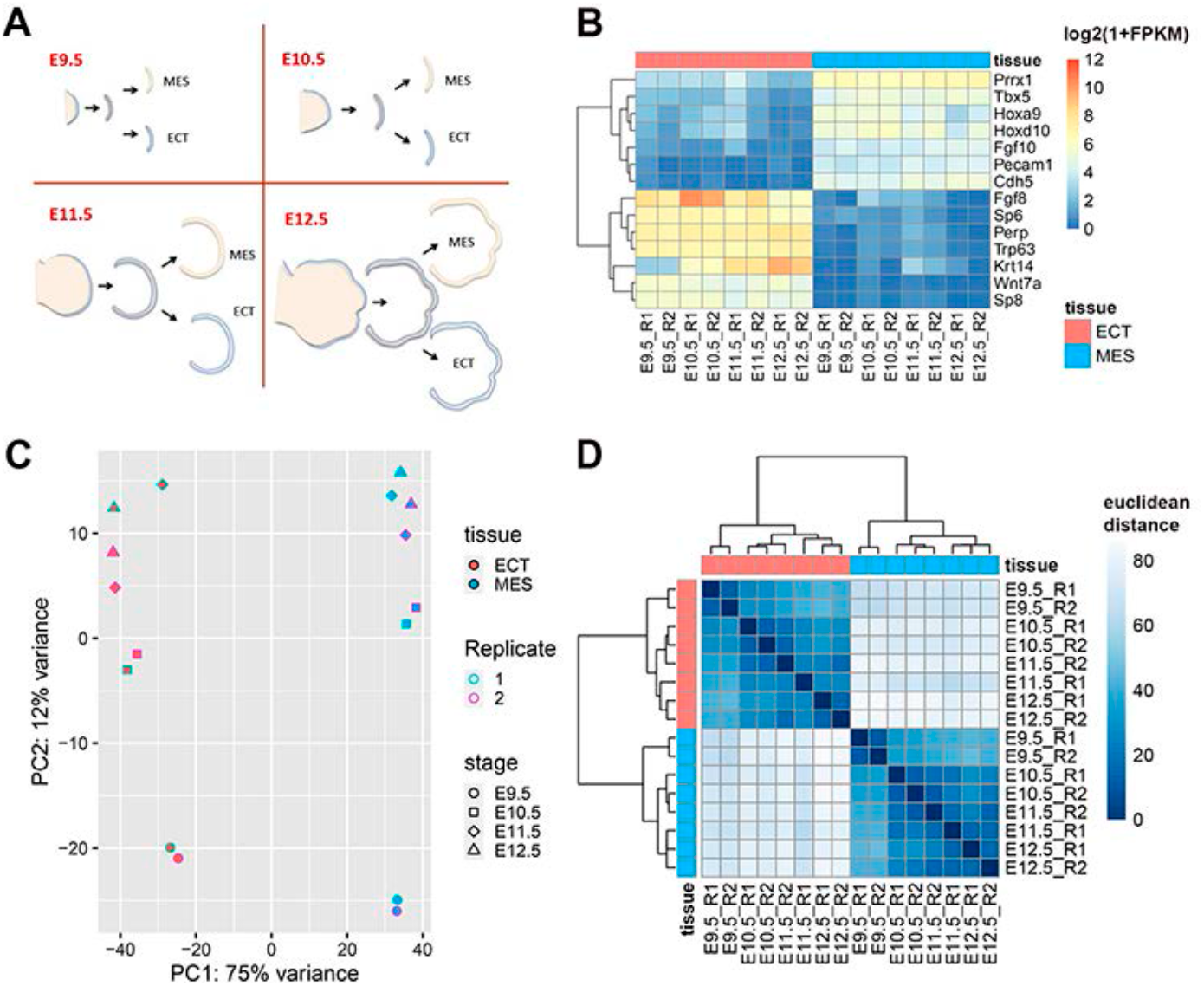
Temporal series of transcriptomes of limb progenitor cells. **A)** Schematic representation of the procedure followed to separate the distal mesoderm from the ectodermal jacket **B)** Heatmap with log2(1+FPKM) values for selected mesodermal (top cluster) and ectodermal (bottom cluster) specific expressed genes, in all samples. **C)** Principal component analysis (PCA) plot of the top 500 most variable genes, using log2(1+FPKM) values. PC1 separates the samples by tissue types and PC2 separates by developmental stage. **D)** The correlation matrix of the top 500 most variable genes, using log2(1+FPKM) values, performed with Euclidean distance and hierarchical ward clustering, shows a strong signal of sample clustering by tissue.

**Figure S2.**
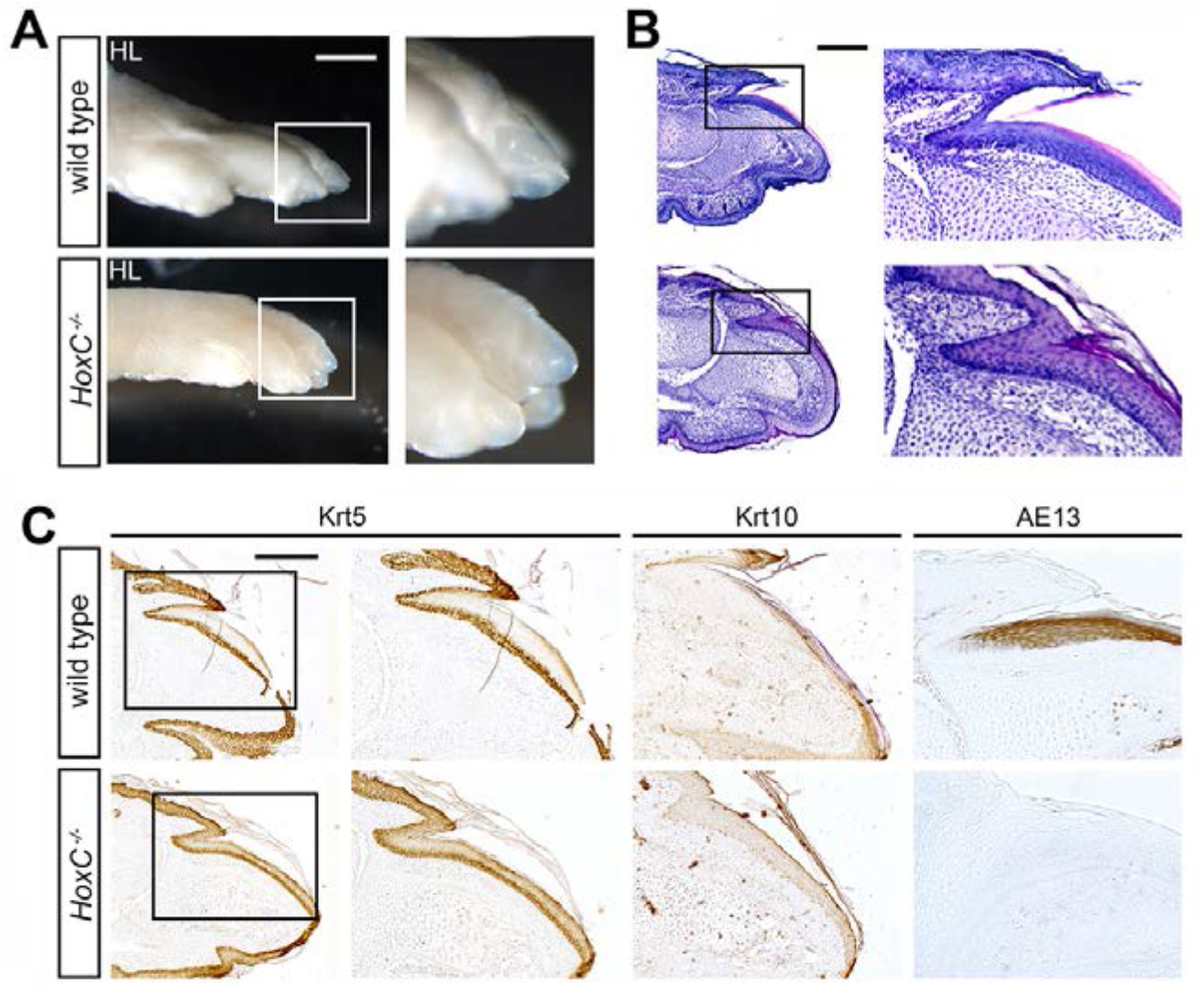
Hypoplastic nails in *HoxC*^−/−^ mutant hindlimbs. **A)** Photographs showing a lateral view of the foot of wild type and *HoxC*^−/−^ newborn mice. Scale bar is 1mm. **B**) Hematoxylin-Eosin stained longitudinal section of a hindlimb digit showing a defective nail organ in the *HoxC*^−/−^ mutant specimen. Scale bar is 200 μm. In **A**) and **B**) the framed area is magnified on the right. **C**) Immunohistochemistry for the detection of Krt5, Krt10 and hard keratins (AE13 antibody) in longitudinal sections of wild type and *HoxC*^−/−^ homozygous newborn mice. For Krt5 immunostaining, a lower magnification is also shown to frame the area under study. Scale bar 200μm. In all sections dorsal is up and distal to the right.

**Figure S3.**
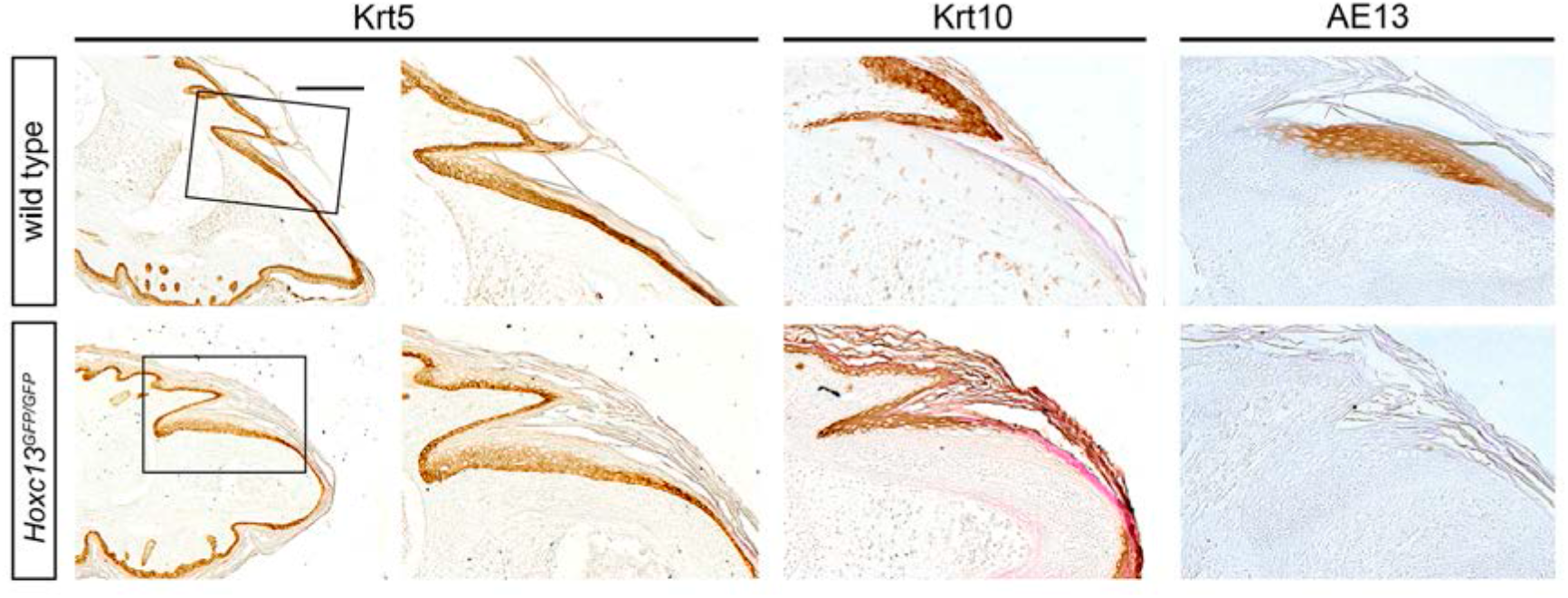
Nail differentiation in *Hoxc13*^*GFP/GFP*^ newborn mice. Immuno-histochemical detection of Krt5, Krt10 and hard keratins (AE13) in longitudinal sections of wild type and *Hoxc13^GFP^* homozygous newborn mice FL digits. The expression of Krt5 was not affected in *Hoxc13*^*GFP*^ homozygous, although Krt10 expression abnormally extended into the supra-basal layer of the nail organ. Note the lack of expression of hard keratins. Scale bar is 200μm.

**Figure S4.**
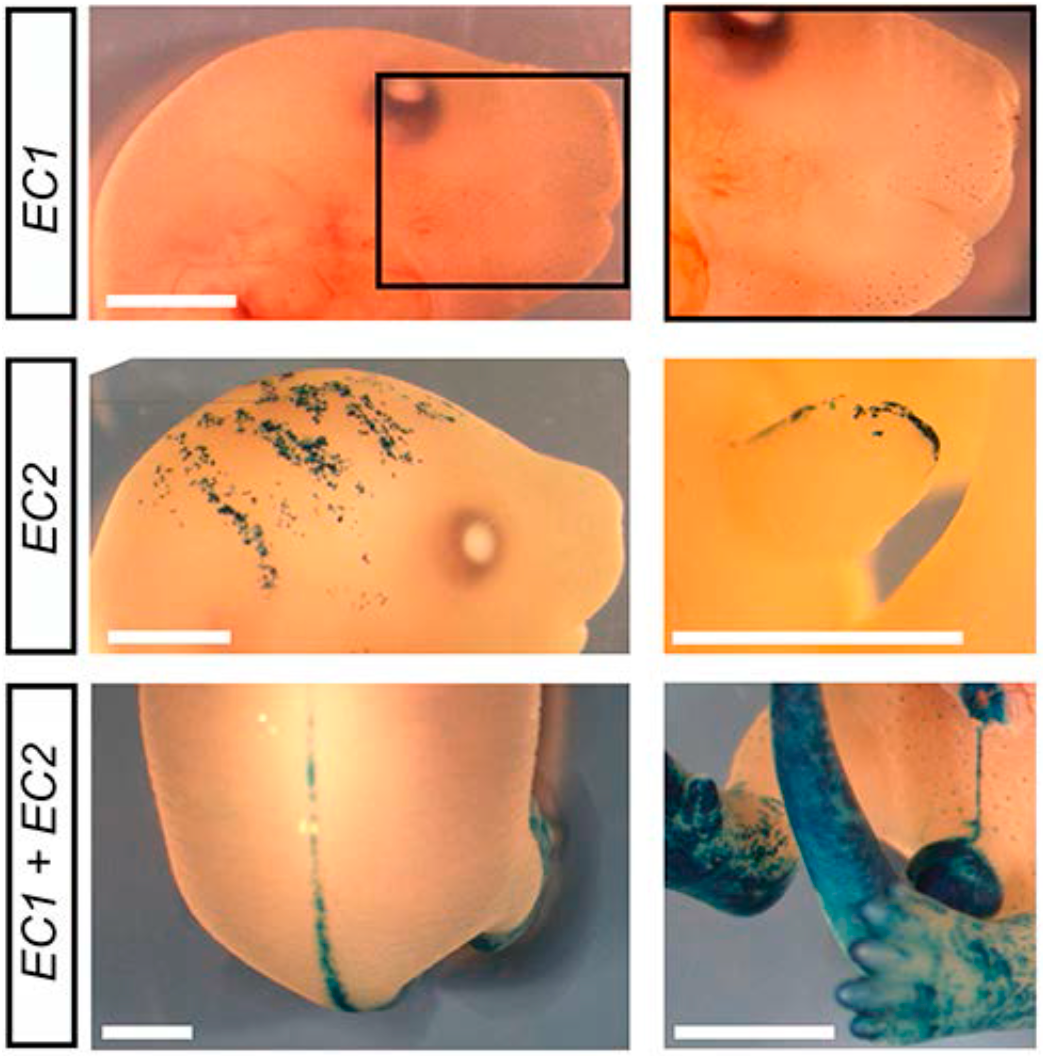
Details of whole-mount *LacZ* pattern driven by EC1 and EC2 putative enhancers. Reporter activity of *EC1*, *EC2* and *EC1*+*EC2* in E16.5 transgenic fetuses. *EC1* showed reporter activity in facial HFs, but not in the whisker pad. *EC2* showed reporter activity in areas of the head ectoderm in variable patterns (left panel) and in the genital tubercle (right panel). *EC1* and *EC2* cloned in tandem showed reporter activity in the dorsal midline (left panel) and in the ventral midline and genital tubercle (right panel). Scale bars 2mm.

**Figure S5.**
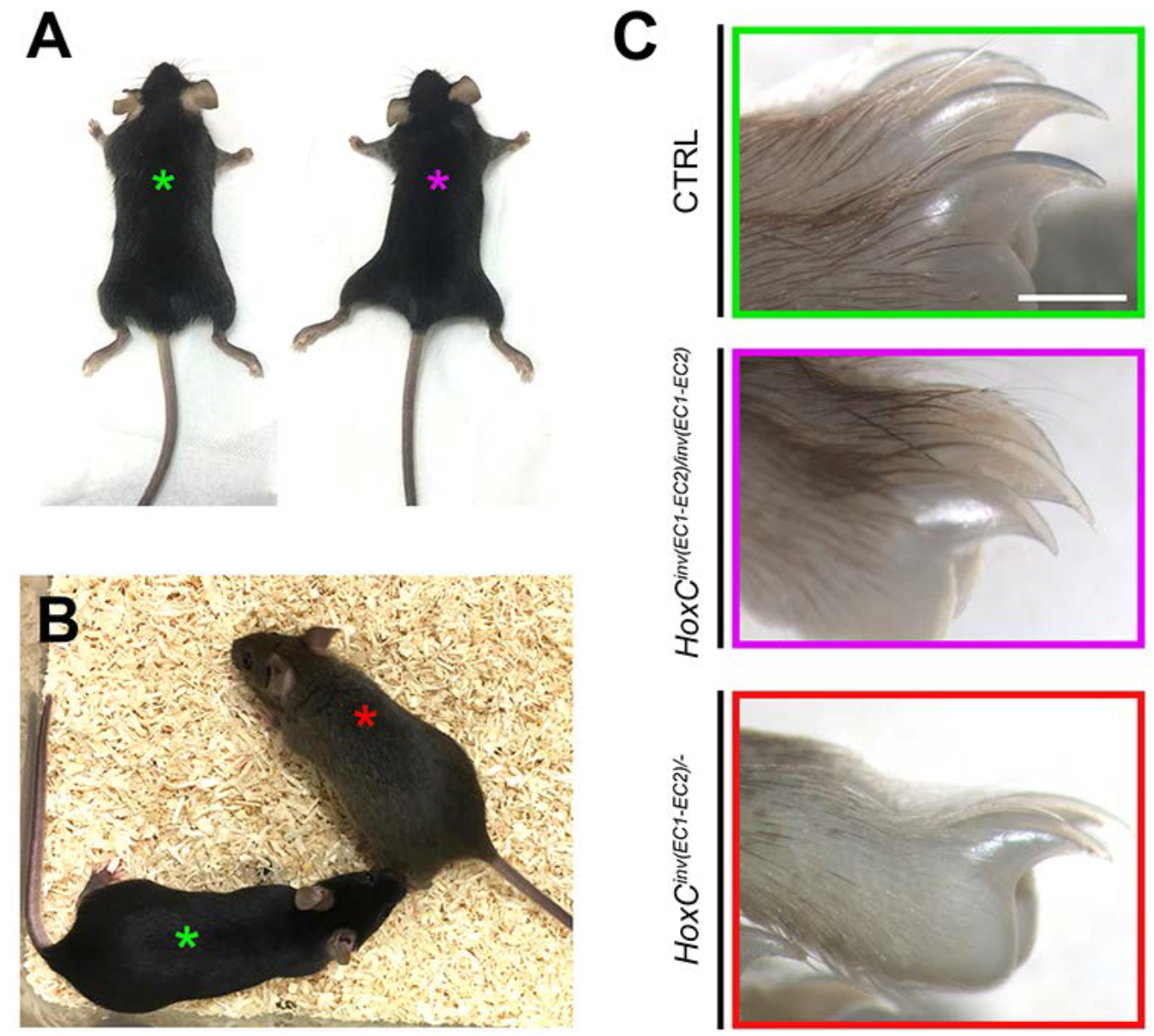
Inversion of the enhancer region. **A)** *HoxC*^*inv(EC1-EC2)*^ homozygous mice (magenta asterisk) were phenotypically indistinguishable from control littermates (green asterisk) and showed wild type nail and hair development. **B)** *HoxC*^*inv(EC1-EC2)*/−^ trans-heterozygous mice (red asterisk) also displayed a normal aspect. **C)** Details of the adult (3 months) hindlimb nails.

**Figure S6.**
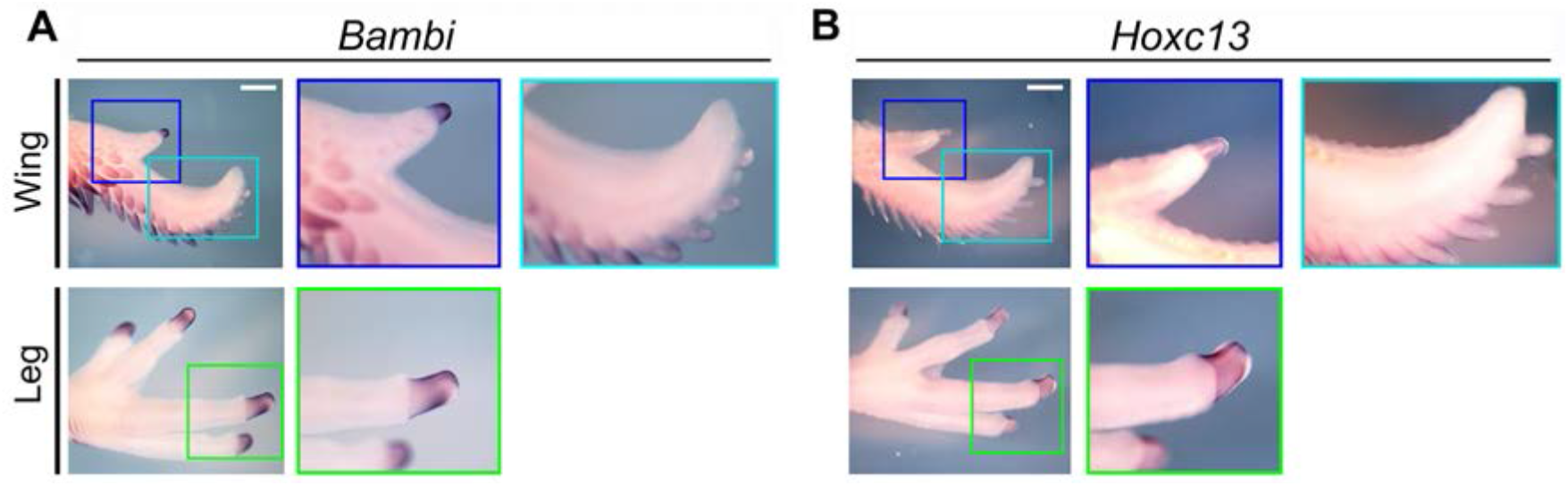
Expression of *Hoxc13* in the chick digit tips. *In situ* hybridization for *Bambi* (**A**) and *Hoxc13* (**B**) in chick digits at Hamburger and Hamilton stage 36 (10 days of incubation). *Bambi* is shown for comparison as a marker of the ectoderm. In the wing, *Hoxc13* is expressed at the tip of digit 1 (blue frame) but is absent from the tips of digits 2 and 3 (cyan frame), which do not form a claw. In the leg, however, *Hoxc13* expression is observed in the tip of all toes; magnified for digits 3 and 4 (green frame).

**Figure S7.**
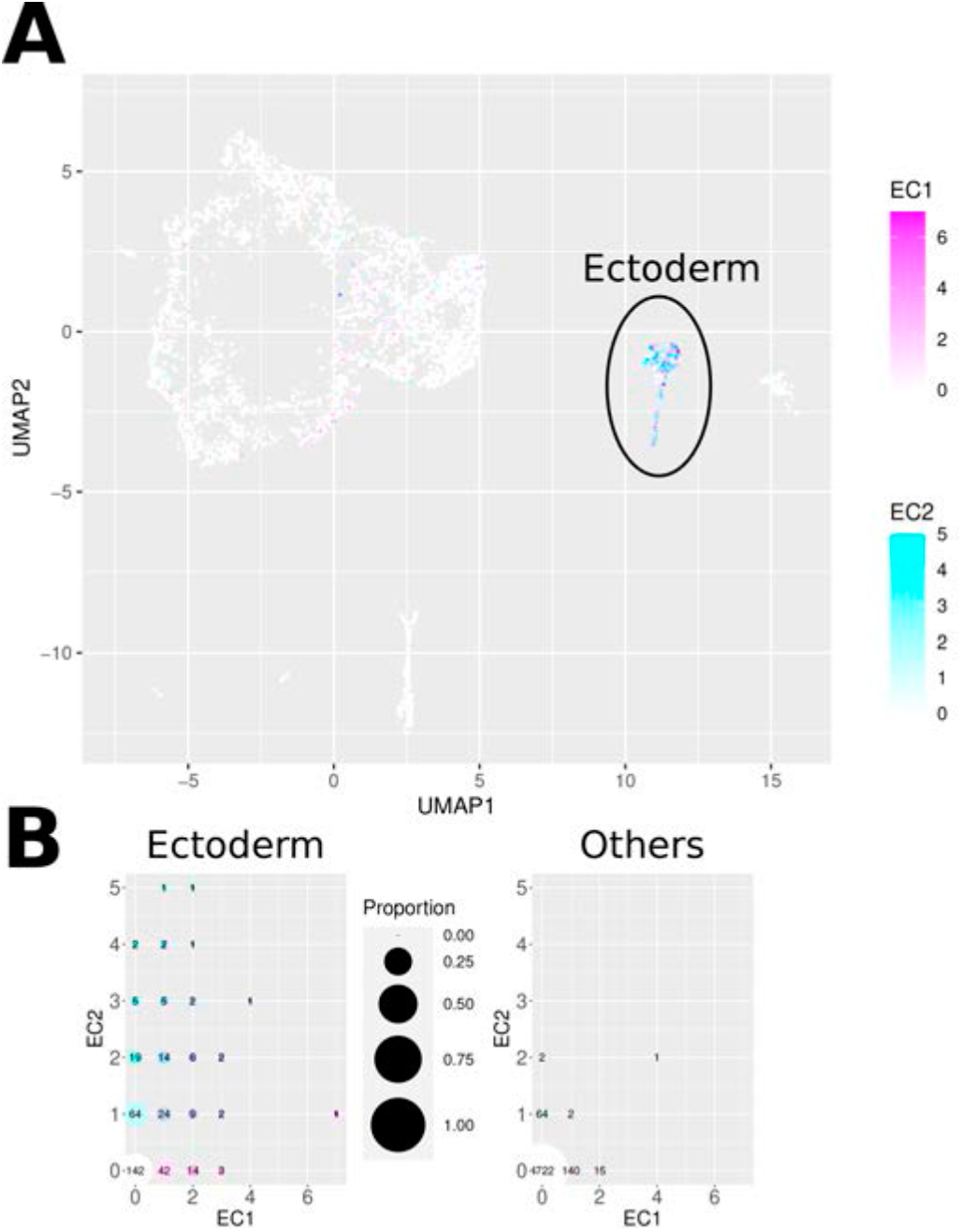
Analysis of the accessibility profile of *EC1* and *EC2* in limb bud scATAC-seq data. **A)** UMAP from Desanlis *et al*. 2020 (10) of single-cell ATAC-seq of control forelimb buds at E11.5 colored by the number of fragments in *EC1* and *EC2*. *EC1* and *EC2* are mainly accessible in the ectoderm cluster. **B)** Number of cells in the ectoderm cluster on the left and on any other cluster on the right for each combination of number of fragment in *EC1* and *EC2*, colored with the same color code as in **A**. In the ectoderm cluster, many cells have both the *EC1* and *EC2* sequences accessible.

## Supplementary Tables

**Table S1.**
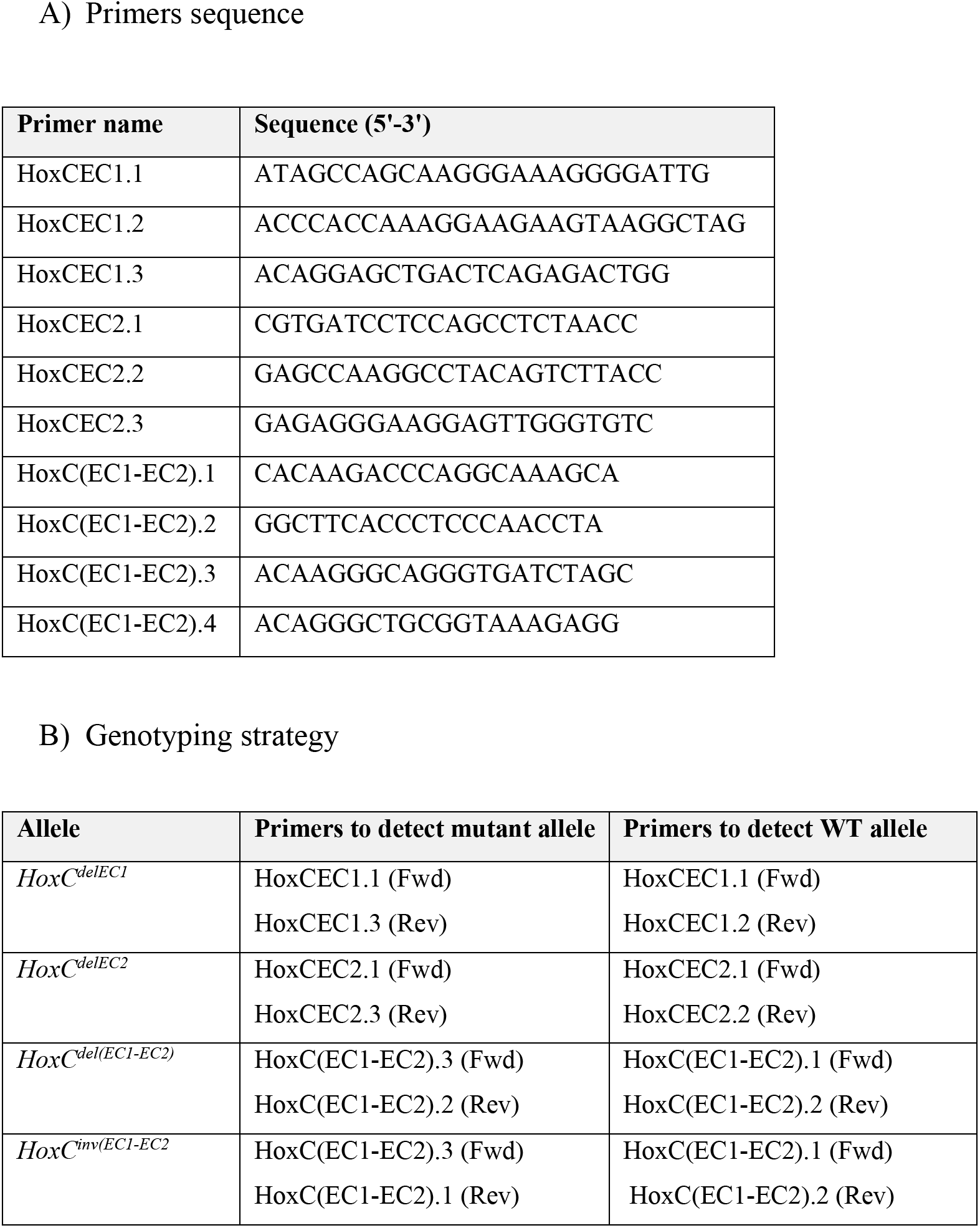
Primers used to genotype *HoxC*^*delEC1*^, *HoxC*^*delEC2*^, *HoxC*^*del(EC1-EC2)*^ and *HoxC*^*inv(EC1-EC2)*^

**Table S2.**
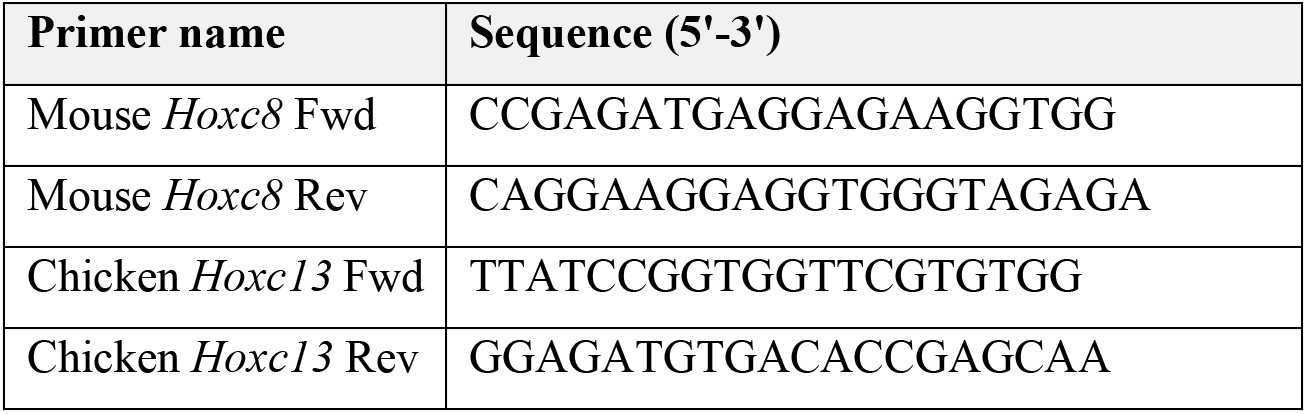
Primers used to amplify cDNA templates for riboprobe synthesis

**Table S3.**
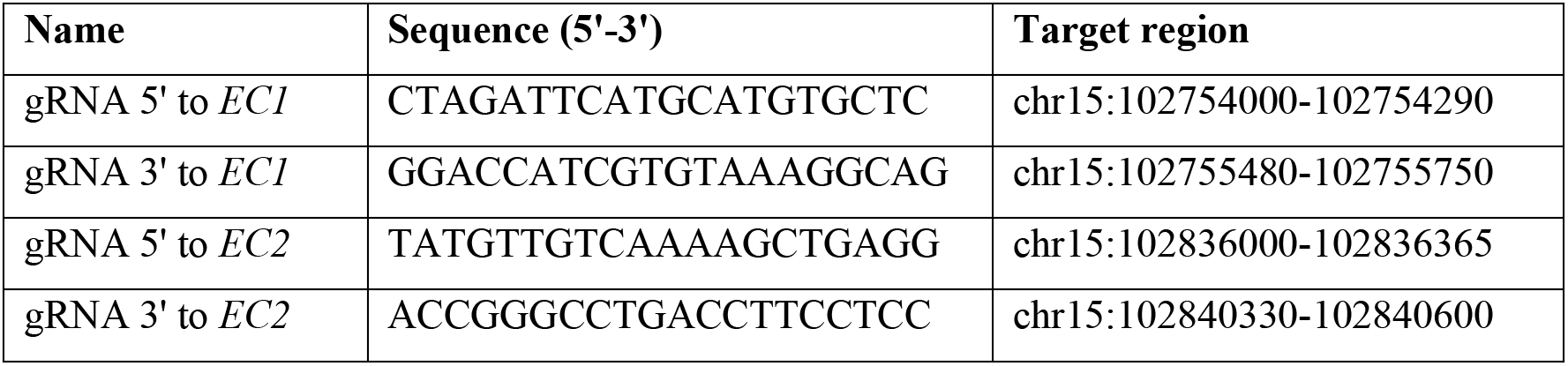
gRNAs used to generate *HoxC*^*delEC1*^, *HoxC*^*delEC2*^, *HoxC*^*del(EC1-EC2)*^ and *HoxC*^*inv(EC1-EC2)*^

